# Development of Retinotopic Feedback: Layer 6 Pyramids to Lateral Geniculate Principal Cells

**DOI:** 10.1101/2024.05.07.592947

**Authors:** William B. Levy, Robert A. Baxter

## Abstract

The development of many feedforward pathways in the brain, from sensory inputs to neocortex, have been studied and modeled extensively, but the development of feedback connections, which tend to occur after the development of feedforward pathways, have received less attention. The abundance of feedback connections within neocortex and between neocortex and thalamus suggests that understanding feedback connections is crucial to understanding connectivity and signal processing in the brain. It is well known that many neural layers are arranged topologically with respect to sensory input, and many neural models impose a symmetry of connections between layers, commonly referred to as reciprocal connectivity. Here, we are concerned with how such reciprocal, feedback connections develop so that the topology of the sensory input is preserved. We focus on feedback connections from layer 6 of visual area V1 to primary neurons in the Lateral Geniculate Nucleus (LGN). The proposed model is based on the hypothesis that feedback connections from V1-L6 to LGN use voltage-activated T-channels to appropriately establish and modify synapses in spite of unavoidable temporal delays. We also hypothesize that developmental spindling relates to synaptogenesis and memory consolidation.

## Introduction

There seems to be as many, or more, positive feedback connections as positive feedforward connections within neocortex and between neocortex and thalamus (***Sherman, 2006***). Thus, we are just as motivated to understand feedback development as to understand feedforward development. During development, e.g. in the visual system, feedforward development, retinal ganglion cells (RGCs) to lateral geniculate nucleus (LGN) principal cells and then LGN to the primary visual cortex (V1), precedes feedback development, V1 layer 6 (denoted by V1-L6 or simply by L6) to LGN. Here we focus on the biological requirements of feedback as opposed to feedforward development. Such biology includes biologically interpreted synaptic modification rules and unavoidable temporal delay aspects of the feedback system. Synaptic modification rules include (i) the activity requirements for synaptogenesis, (ii) a spike-timing rule for associative strengthening versus disassociative weakening of an existing synapse, and (iii) a rule for discarding a sufficiently weakened, existing synapse.

We insist that synaptic modification rules must be biologically plausible and must lead to stable, steady-state values (convergence) of connectivity in stationary, ergodic input environments. Because most of the connections of neocortex and corticothalamic systems are excitatory, we conclude they encode positive correlations. Convergence of positively correlated inputs creates neural transformations that can be used predictively, and that can be viewed as compressive. Convergence of inputs that contain a positive correlation in a feedforward network are of two types. If the neuron is class-supervised, the positive correlation is between the feature and the class the neuron represents. If the neuron does not receive a class signal but only receives feature signals, then the positive correlations are between the features themselves, and we call such neurons unsupervised (more expressively, they could have been called self-supervised). The set of acquired synapses determines a neuron’s best stimulus and, at least in this sense, the identity of the neuron both in theoretical terms and in the classic perspective of neurophysiologists Hubel and Wiesel (***Hubel and Wiesel, 1962, 1963***). Technically, from a probabilistic perspective, this best stimulus of neurophysiology is a neuron’s latent variable. An example of such a positive correlation-based latent variable is retinotopy.

Retinotopy refers to the spatial organization of neurons that reflects the neighborliness of the retinal input; in other words, retinotopy represents the spatial mapping of visual input from the retina to neurons. Retinotopy develops via topologically induced positive correlations among nearby neurons. Successive brain regions in the visual system reproduce, to some extent, such retinotopy. When feedforward and feedback pathways exist, retinotopy of connectivity is usually assumed to exist in both the feedforward and feedback directions. In neuroscience, this is called “reciprocal” connectivity, or to avoid certain ambiguities in the jargon, “in-register” reciprocity. For example, the topology of V1-L6 projections into the LGN will tend to exhibit nearly identical retinotopy as the retinotopy of the LGN projections to V1. Where this in-register reciprocal connectivity has been carefully looked for, there is evidence for a certain amount of maintained topology of feedback connections in early sensory processing (***Sikkens et al., 2019; Ren et al., 2019***). Indeed, topological feedback is often assumed in models of attention where positive feedback can amplify particular subsets of neuron firings. The neural anatomical idea of in-register reciprocity for feedforward-feedback systems is anticipated within the machine learning literature by the symmetry of synaptic weights, *w*_*ij*_ = *w*_*ji*_, fundamental to the networks of ***Hopfield*** (***1982***) and of ***Cohen and Grossberg*** (***1983***).

Here, we focus on the special requirements of feedback development, compared to feedforward development, necessary to produce feedback retinotopy. This focus includes a proposed biophysical mechanism that distinguishes Nature’s implementations of feedforward development using the NMDA receptor versus feedback development using the voltage-activated T-channel. Here, “T-channel” refers to a T-type Ca^++^ channel. The proposed model provides a spindle hypothesis that includes an explanation of how the T-channel solves timing and delay issues in the development of feedback connectivity.

Although the initiating event of an adult spindle is controversial, here we formulate a spindle hypothesis based on ***Grenier et al. (1998***) and ***Llinás and Steriade*** (***2006***), given that this unknown, spindle-initiating event has occurred. To our mind, the critical biophysics of developmental spindling is selective principal cell activation by the retina upon the LGN, which leads to specific hyperpolarization of principal cells in the LGN. This cell-specific hyperpolarization not only halts the RGC-initiated firing but de-inactivates (i.e., primes) T-channels in the dendrites of just those principal cells that were selectively fired. The cell-specific de-inactivation of T-channels creates the condition that only those particular principal cells burst-fire when receiving feedback synaptic activation. During development, we assume hyperpolarization is selective to the LGN principal cells recently activated by spontaneous RGC activity caused by retinal waves. This selectivity is critical to the model as it creates a topographically specific and brief memory of the original retinal activation.

Under a spike-timing assumption that requires presynaptic activation (i.e., L6 excitation) to precede postsynaptic activation (i.e., LGN excitation), we come upon one possible explanation for the T-channel and its unusual priming requirement: the requirement that allows low-voltage excitation to produce burst firing and a large calcium influx. This large calcium influx provides the control mechanism for spike-timing of associative synaptic potentiation and synaptic depression in feedback pathways, in contrast to control via NMDA receptors in feedforward systems. The initial excitation, RGC to LGN, comes first, but its memory (via cell-specific hyperpolarization) preserves the pre-then-postsynaptic rule (***Llinás and Steriade, 2006; Payeur et al., 2021***) for the loop-delayed arrival of L6 (presynaptic) activation and the follow-on T-channel activation (i.e., de-inactivation plus synaptic activation). Note that as a T-channel is calcium conducting, large transient increases of postsynaptic, intracellular Ca^++^ on a dendrite is the signal for increasing synaptic efficacy, much like NMDA receptor associative potentiation.

Putting these ideas into equations, we elucidate a dynamic model of retinotopic development for feedback. With the strong suggestion of the well-supported correlations between retinally-driven V1 spindles and such feedback development (***Murata and Colonnese, 2016***), the model solves the spike-timing problem posed by feedback development. Specific RGC activity associated with retinal waves activates, in a cell-specific way, LGN principal cells. In the hypothesis, it is only at these cells that a hyperpolarization occurs. This hyperpolarization (i) replaces the usual spike-timing synaptic potentiation/depression rules (***Levy and Steward, 1979, 1983***), (ii) serves as a short-term memory marking the recently activated LGN primary neurons, and (iii) primes the local T-channels into de-inactivation. The repetitive nature within each spindle permits synaptogenesis and synaptic potentiation on synapses associated with recent RGC activity (see dynamic equations 8 and 10 in Model).

## Model

### Model Assumptions

The model consists of three neural layers: the RGC layer, the LGN layer, and the L6 layer. The RGC layer is consists of a linear array of neurons (1-D) and opponent coding is ignored.

The developmental model here assumes certain processes precede adaptive synaptogenesis. While the model does not require existing aligned connections from L6 to the LGN, in our demonstrations here we chose to start with a mixture of a small number of connections, some of which are aligned. These connections are assumed to be the result of neurotropic chemotaxis.

In general, both feedforward connections from LGN to L6 and feedback connections from L6 to LGN develop via adaptive synaptogenesis. These connections can be modified in three ways: formation of synaptic connections via synaptogenesis, modification of their synaptic strengths, and shedding of connections. Feedforward connections from LGN to L6 are associated with a two-layer, developmental, feedforward, adaptive synaptogenesis model. The feedforward connections (and synaptic strengths) are assumed to develop and stabilize prior to the development of feedback connections. The separation of the development of the feedforward and feedback connections need not be strict, but in the simulations presented here we assume there is no overlap in the development of the feedforward and feedback connections. With the feedforward model, connections from LGN to L6 can be shown to develop retinotopic connectivity. Feedforward connections are known to be topographic by PN8, in particular RGC to primary thalamocortical (TC) neurons and the L6 recipients of these TC inputs. In the simulations presented here, the feedforward connections from LGN to L6 are assumed to be retinotopic and fixed in order to focus on the development of the feedback connections.

Retinal inputs consist of *n*_*a*_ active, adjacent RGC neurons firing for a prescribed duration, *τ*_*R*_. Each retinal input pattern is specified by a center location on the retina, *i*_*c*_, and the number of active retinal inputs, *n*_*a*_. The animal is assumed to dwell on each different retinal input pattern for a duration *τ*_*d*_. Once the RGC excites an LGN neuron, and subsequently an aligned L6 neuron, connections from L6 to the LGN begin to form. When a sufficient number of connections from L6 to the LGN are created, feedback oscillations will tend to continue regardless of the RGC input pattern duration. In the simulations, the feedback oscillations cease when the dwell time is exceeded. The time between spindle events (groups of bursts associated with successive feedback oscillations), which may actually last for several seconds *in vivo*, is limited to a few time steps in the simulations. When the dwell time *τ*_*d*_ is exceeded, all activities are reset and the next retinal input pattern is presented. Of course, the model could be made more realistic by inserting a longer inactive period between spindle events, perhaps with some amount of random deviation from a specified duration of inactivity.

An LGN primary neuron will generate an output (initial segment) spike if (a) the neuron is at rest and the RGC input is above a threshold, or (b) a T-channel spike occurs. The LGN spike produces dendritic hyperpolarization, and this hyperpolarization primes (inactivates) the T-channel. A primed T-channel can easily de-inactivate and spike with just a few mV rise in excitation. T-channels are excited by L6 input to the LGN.

For feedback connections, both synaptogenesis and synaptic modification require recent hyperpolarization. Synaptogenesis occurs when the T-channel is not spiking and synaptic modification occurs when the T-channel is spiking.

Spindle events in the neocortex reflect cyclical activations of selected LGN primary neurons. In the visual cortex prior to eye-opening, a single spindle event consists of 8 to 14 cycles repeated at 15 to 30 Hz, whereas in the adult the frequency range is 8 to 15 Hz (***Clawson et al., 2016; Llinás and Steriade, 2006***). The number of spindle oscillations depends on the strength of the positive feedback loop. This feedback loop requires a minimum amount of symmetry between the LGN to L6 connections and the L6 to LGN connections. Here, parameters are typically set such that feedback cycles repeat at a frequency of 20 Hz (corresponding to a 50 msec delay), there are 9 bursts per spindle event with one spike per burst, and there is one spindle event for each retinal input pattern.

A diagram of the model is shown in Figure 1.

**Figure 1.**
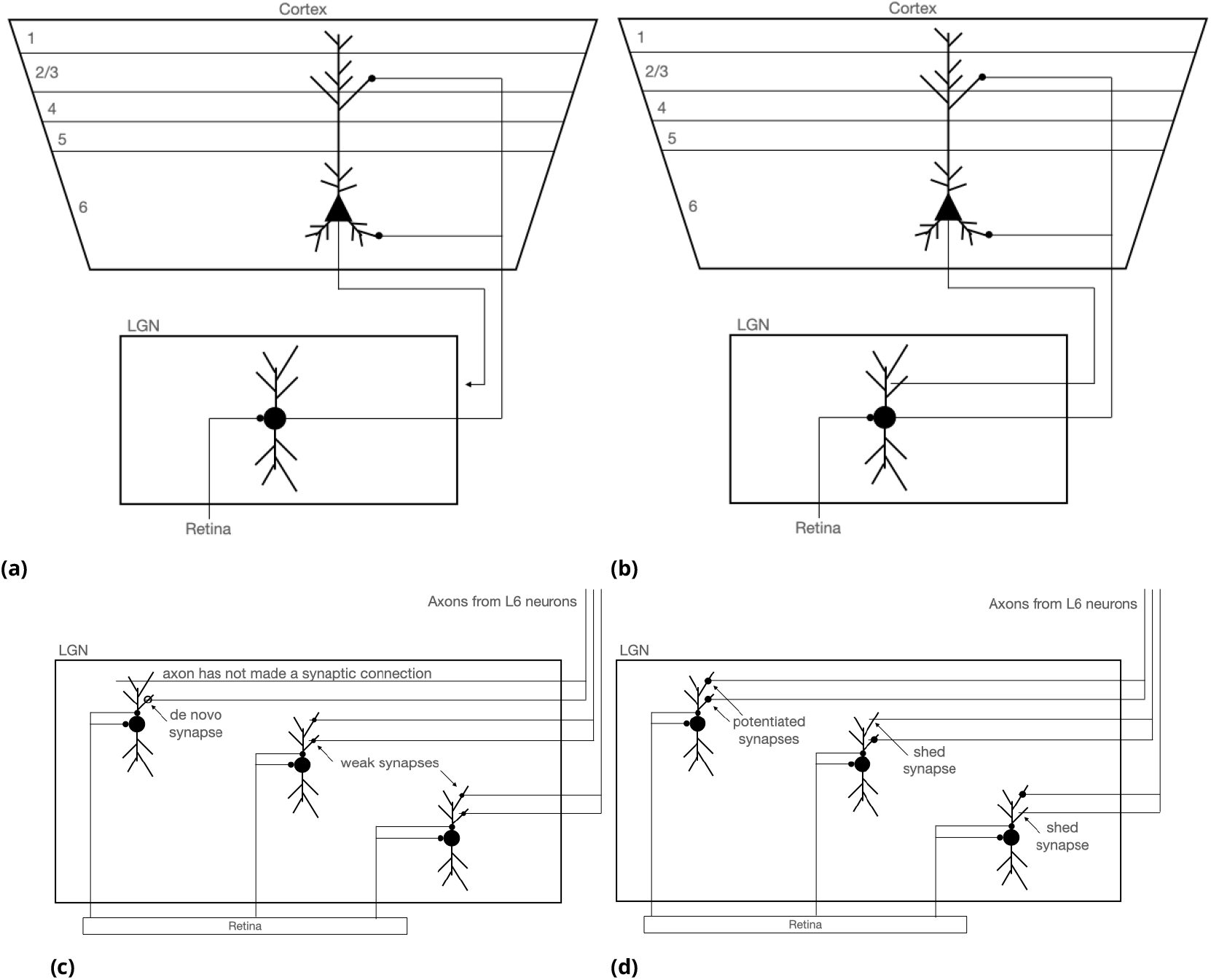
Visualization of L6 to LGN synaptic development for LGN principal neurons. (a) The initial state of most L6 axons is just outside of the LGN. (b) Around PN5 through PN7, axons begin to invade LGN (***Seabrook et al., 2013***). Here, initial connectivity is developed as a biased random process. The biasing reflects the chemotactic signaling of retinotopy. (c) A mismatch between retinal activity-induced hyperpolarization/de-inactivation and T-channel activation on any one neuron raises the probability of synapse formation for that neuron. (d) De novo synapses, having been formed by either of the random mechanisms (b) or (c), are now modified via equation (8). This equation can strengthen (LTP) or weaken (LTD) these de novo synapses. If sufficient LTD occurs, such a synapse is discarded (shed).

### Notation, Neuronal excitations and outputs, and Delays

Herein, the word “aligned” is meant to denote the retinotopic alignment between layers. Only aligned LGN neurons undergo hyperpolarization and T-channel spiking.

In the equations, RGC neurons are indexed by *i*, LGN neurons are indexed by *j*, and L6 neurons are indexed by *k*. RGC neurons represent the retinal input and their activities are denoted by *x*_*i*_(*t*). LGN neuronal excitation by RGC neurons is denoted by *y*_*j,R*_(*t*); excitation of the T-channel is denoted by *y*_*j,T*_ (*t*). An LGN spike caused by retinal input is denoted by *z*_*j,R*_(*t*), an LGN initial segment (IS) spike is denoted by *z*_*j,IS*_ (*t*), whereas a T-channel spike is denoted by *z*_*j,T*_ (*t*). Neuronal excitations in L6 are denoted by *y*_*k*_(*t*) and L6 output spikes are denoted by *z*_*k*_(*t*). Spikes are binary; i.e., *z*_*j,R*_(*t*), *z*_*j,IS*_ (*t*), *z*_*j,T*_ (*t*), and *z*_*k*_(*t*) are restricted to 0 or 1.

An LGN neuron becomes hyperpolarized whenever an initial segment spike is produced. The hyperpolarization indicator function, *h*_*j*_ (*t*) ∈ {0, 1}, turns on whenever *z*_*j,IS*_ (*t* − 1) = 0 and *z*_*j,IS*_ (*t*) = 1 and turns off after a delay of *τ*_*h*_. In the simulations presented here, the hyperpolarization duration is set equal to the dwell time.

The synaptic connections and strengths (weights) of the connections from L6 to LGN are denoted by *c*_*kj*_ (*t*) and *w*_*kj*_ (*t*), respectively, where the *c*_*kj*_ (*t*) are 0 or 1 and the *w*_*kj*_ (*t*) are nonnegative real values. When graphing the weights as images or as a function of *k* or *j*, we assume that neurons *k* and *j* are equally spaced. This assumption is not a limitation of the model; given the distances between neurons *k* and *j*, however nonlinear, the graphs can use the distances provided. Lacking the distances between neurons, we make the simplifying assumption that the neurons are equally spaced.

The following delay parameters are included in the three-layer model: *τ*_12_ is the delay from the RGC layer to the LGN layer (5 msec), *τ*_23_ is the delay from LGN to L6 (25 msec), and *τ*_32_ is the delay from L6 to LGN (25 msec). The time it takes a signal to propagate from the LGN to L6 and back to LGN is denoted by *τ*_232_ = *τ*_23_ + *τ*_32_ (50 msec).

### Spindling Event Sequence

In the simulations, LGN spindling is assumed to consist of the following sequence of events :

1. Initial inactivity
2. Onset of next retinal input pattern and activation of the RGC layer
3. Bursting caused by RGC input or T-channel firing
  a. Initial segment firing caused by RGC input to the LGN or by T-channel firing
  b. Local hyperpolarization of aligned LGN neurons
  c. Burst: sequence of spikes by hyperpolarized LGN neurons when L6 input is sufficient
  d. Synaptic modification (and shedding) occurs after each spike in the burst
  e. Synaptogenesis occurs whenever the LGN neuron is hyperpolarized and the T-channel is not firing
  f. Inactivity
4. Repeat until all retinal input patterns have been presented *n*_*p*_ times

In the simulations, the timing of the spikes in a burst firing sequence is specified by the number of spikes per burst, *n*_*s*_, and the interspike interval (ISI) between spikes in a burst, *τ*_*ISI*_. Also, a refractory period of inactivity, with duration *τ*_*IA*_, occurs at the end of each burst. For example, *n*_*s*_ = 2 and *τ*_*ISI*_ = 4 msec specifies a burst of two spikes separated by 4 msec which implies a burst frequency of 250 Hz.

Note that in order to permit T-channel firing and maintain hyperpolarization, the duration of inactivity after a burst must be less than the feedback delay and the duration of hyperpolarization must be greater than the feedback delay, i.e. *τ*_*IA*_ < *τ*_23_ + *τ*_32_ < *τ*_*h*_.

### LGN Layer Excitations and Outputs

The excitation of LGN neuron *j* by the RGC is the sum of its retinal inputs,

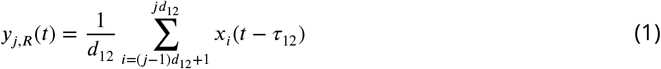

where *d*_12_ is the RGC-to-LGN connectivity decimation factor. Here, we set *d*_12_ = 1 which implies *y*_*j,R*_(*t*) = *x*_*i*_(*t* − *τ*_12_) only when *j* = *i* and 0 otherwise (perfect alignment from RGC to LGN). The excitation of LGN neuron *j*’s T-channel by L6 is the sum of its L6 inputs,

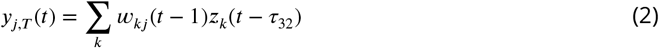

If the LGN neuron was previously at rest and the current excitation by the RGC is above the threshold, then the RGC input will induce an IS spike. This RGC-induced spike is denoted by *z*_*j,R*_(*t*) and can be expressed as

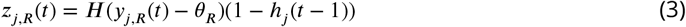

where *θ*_*R*_ is the firing threshold for RGC-induced spikes, and *H*(·) is the unit step function defined as *H*(*x*) = 1 if *x* > 0 and 0 otherwise.

A T-channel spike, denoted by *z*_*j,T*_ (*t*), will be produced when the T-channel excitation exceeds a firing threshold and the LGN neuron *j* is hyperpolarized, i.e.

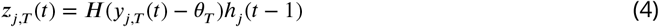

where *θ*_*T*_ is the firing threshold for the T-channel spike.

A T-channel spike will produce an IS spike regardless of *y*_*j,R*_(*t*), and this IS spike will hyperpolarize the dendrite so that *y*_*j,R*_(*t*) = *y*_*h*_. Therefore, *z*_*j,R*_(*t*) and *z*_*j,T*_ (*t*) are mutually exclusive and equation 3 can be combined with equation 4 to produce the following expression for an LGN IS spike,

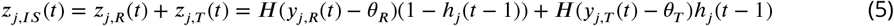

### L6 Layer Excitations and Outputs

Neurons in the L6 layer are excited according to

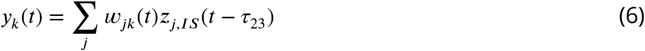

where the *w*_*jk*_(*t*) are the feedforward LGN→L6 synaptic weights. For the simulations described in the Results, we have assumed that the feedforward weights are fixed and produce perfect retinotopic connectivity from LGN to L6; i.e., we have assumed that the number of neurons in the LGN and L6 are equal and that the feedforward weights provide perfect alignment from LGN to L6.

L6 neurons emit a spike when *y*_*k*_(*t*) is greater than a threshold,

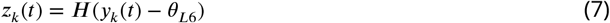

where *θ*_*L*6_ is the firing threshold for L6 neuronal spikes.

### Synaptic modification and shedding

Synaptic modification and shedding occurs at every time step. In the Hebbian spike-timing potentiation/depression rule, the presynaptic signal is modeled as the activation of an NMDA-receptor. This receptor has a slight delayed onset of the required glutamate binding and a very slow offset of unbinding. The presynaptic signal is modeled as a saturate and decay mechanism driven by the L6 output, *z*_*k*_(*t*), with biologically appropriate rise and decay times. This presynaptic signal is referred to as “z bar” and denoted with a tilde, 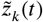. The rise time to peak (saturation) is 10 msec and the decay time from peak to 1/*e* is 100 msec. In fact, this is a deterministic representation of a stochastic process; i.e., the average availability of a glutamate-primed NMDA-receptor awaiting postsynaptic depolarization for activation. The equation that embodies this molecular basis of pre- and post-synaptic associative modification is given by

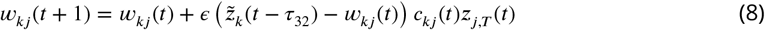

The presynaptic signal 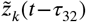 behaves much like a capacitor which is either charging to saturation or discharging. The rate of synaptic modification is governed by *ϵ*. The term *c*_*kj*_ (*t*) ensures that only existing connections are able to modify their synaptic strengths. The term *z*_*j,T*_ (*t*) requires T-channel firing for synaptic modification to occur.

For stationary, ergodic retinal input, equation 8 leads to synaptic weights that approximate the conditional expectation 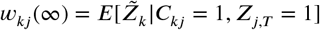 where the upper case variables represent the corresponding random variables associated with the lower case variables.

Synaptic shedding (synapse removal) occurs when *w*_*kj*_ (*t*) < *θ*_*w*_ for any *t* (typically, *θ*_*w*_ = 0.03), i.e.

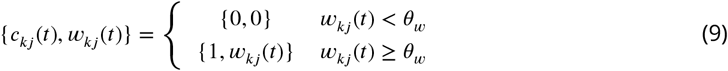

### Synaptogenesis

Synaptogenesis occurs whenever an LGN neuron is hyperpolarized and the T-channel is not firing. Synaptogenesis of an L6 to LGN synapse is a conditional Bernoulli process for each possible (*k, j*) pair. Duplicate synapses are not permitted. The probability of forming a synapse between L6 neuron *k* and LGN neuron *j* is

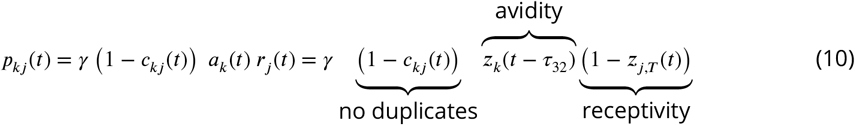

where *γ* is a positive synapse formation rate factor, *a*_*k*_(*t*) is the presynaptic avidity, and *r*_*j*_ (*t*) is the postsynaptic receptivity. The term 1−*c*_*kj*_ (*t*) prevents duplicate synapses. In all simulations here, the avidity factor is set to the L6 neuron output, i.e. *a*_*k*_(*t*) = *z*_*k*_(*t* − *τ*_32_). The postsynaptic receptivity term 1 − *z*_*j,T*_ (*t*) prevents synaptogenesis when the T-channel is firing. In words, a non-zero probability of forming a synapse from L6 to LGN requires three tests be satisfied:

Test 1: a connection from L6 neuron *k* to LGN neuron *j* does not exist (*c*_*kj*_ (*t*) = 0)

Test 2: the L6 neuron is firing (*a*_*k*_(*t*) = 1)

Test 3: the LGN T-channel is not firing (*z*_*j,T*_ (*t*) = 0)

The parameter *γ* governs the rate of synaptogenesis. New connections have a synaptic weight of *W*_0_ (0.1 in all of the simulations presented here).

Equation 10 is referred to as *restricted* or *spatially-biased* synaptogenesis, and it produces the most efficient synapse formation. In contrast, *unrestricted synaptogenesis* does not make use of the avidity or receptivity terms and is independent of hyperpolarization and T-channel firing. The equation for unrestricted synaptogenesis simplifies to

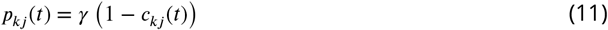

which implies that the probability of synapse formation is a constant for nonexistent synapses.

The problem with equation (11) is that new synapses are created where no connections exist and if these connections are not aligned with RGC activities they will be shed. That is, connections are created and shed in unaligned regions and synaptogenesis never shuts off.

Therefore, to solve this problem for the unrestricted case, we have examined five additional mechanisms for shutting down synaptogenesis and stabilizing the connectivity:

1. Specify a synaptogenesis shutdown time. This method is the simplest method. It is similar to establishing a critical period for synaptogenesis. (Requires one parameter.)
2. When the average LGN T-channel excitation, 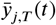, exceeds a threshold, turn off synaptogenesis. In this case, the average T-channel excitation is updated only when instantaneous LGN retinal-based excitation, *y*_*j,R*_(*t*), exceeds the threshold, *θ*_*R*_. Because the activity wave travels across the retina, the average T-channel excitation must only be updated when the retinal-based excitation is sufficiently high. The basic idea is that high average T-channel excitations should lead to reduced probabilities of synapse formation. (Requires two parameters plus the average update rate parameter.)
3. When the sum of the weights exceeds a threshold, turn off synaptogenesis. The rationale for this method is that when the sum of the weights exceeds a threshold, the existing connectivity is sufficient and no additional connections are needed. (Requires one parameter.)
4. Decrease the value of *γ* for a synapse when its synaptic weight exceeds a threshold. This method is similar to that used in (***Levy and Baxter, 2023***). The basic idea is that the probability of forming new synapses should decrease when the weights exceed a specified value. This method assumes that each synapse has a specific value of *γ*. (Requires two parameters plus the value of *γ* for each synapse.)
5. When the spatial distance between the L6 and LGN neurons exceeds a threshold, decrease the probability of synapse formation. This spatial distance restriction method reduces the probability of synapse formation as the spatial distance between the neurons increases. (Requires one paramter.)

In Results, we provide examples of shutting down synaptogenesis for methods 3 and 5.

Adding a weight sum threshold restriction to the unrestricted case (method 3) is expressed as

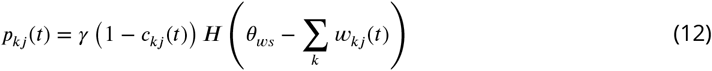

where *θ*_*ws*_ is the weight sum threshold.

Adding a spatial distance restriction to the unrestricted case (method 5) is expressed as

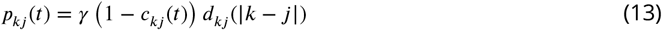

where *d*_*kj*_ (|*k* − *j*|) is a function of the spatial distance between the L6 and LGN neurons with a distance threshold *θ*_*d*_ as a parameter. The distance threshold determines when the distance function decreases to a minimum value *d*_min_; *d*_*kj*_ = *d*_min_ when |*k* − *j*| ≥ *θ*_*d*_. We implemented a variety of distance attenuation functions including a boxcar (1 if less than the threshold and *d*_min_ otherwise), a logistic, a cosine, a parabola, and a generalized Gaussian. In Results, we used a generalized Gaussian function with parameter *β* because it provides a simple and flexible way of controlling the shape of the distance function (see Figure 11). The generalized Gaussian as a spatial distance attenuation function is computed as

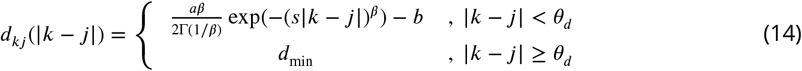

where Γ(·) is the gamma function, *s* is a distance scaling factor (*s* = 0.1), and *β* > 0 is a parameter. The constants *a, b* > 0 scale *d*_*kj*_ (|*k* − *j*|) to the closed interval [1, *d*_min_].

### Synaptic transmission failures

Synaptic transmission failures are common in neocortex. Synaptic failures affect the model in two ways: (a) failure to transmit an L6 neuron output to the LGN, and (b) failure of 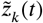 to rise to saturation. Synaptic transmission successes are modeled as Bernouli processes with probability 1 − *f* where *f* is the failure rate. In all simulations here, the synaptic failure rate is set to 0.

### The feedforward model

In the feedfoward model, the excitation of LGN neurons is denoted simply by *y*_*j*_ (*t*) and is identical to *y*_*j,R*_(*t*), *y*_*j*_ (*t*) = *y*_*j,R*_(*t*). The output of LGN neurons in the feedforward model is *z*_*j*_ (*t*) = *H*(*y*_*j*_ (*t*) − *θ*_*R*_). The excitation of neurons in L6 is given by

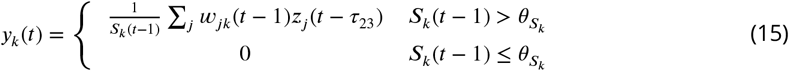

where *S*_*k*_(*t*) =∑ _*k*_ *w*_*jk*_(*t*) and 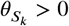. The output of L6 neurons *z*_*k*_(*t*) = *H*(*y*_*k*_(*t*) − *θ*_*L*6_).

Synaptic modification is governed by the equation

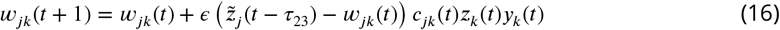

where the presynaptic input is the saturate and decay function related to *z*_*j*_ (*t* − *τ*_23_). Synaptic shedding (synapse removal) is similar to shedding in the feedback model.

Synaptogenesis occurs whenever an L6 neuron is not firing. Synaptogenesis of an LGN to L6 synapse is a conditional Bernoulli process for each possible (*j, k*) pair. Duplicate synapses are not permitted. The probability of forming a synapse between LGN neuron *j* and L6 neuron *k* is

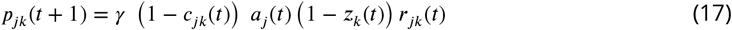

where *γ* is a synapse formation rate factor and *a*_*j*_ (*t*) is an avidity factor. In all simulations here, the avidity factor is set to the delayed LGN neuron output, i.e. *a*_*j*_ (*t*) = *z*_*j*_ (*t* − *τ*_23_). The term 1 − *c*_*jk*_(*t*) prevents duplicate synapses, and the term 1 − *z*_*k*_(*t*) prevents synaptogenesis when the L6 neuron is firing. The term *r*_*jk*_(*t*) is an alignment restriction factor, which is typically related to the running averages of the LGN outputs, e.g. 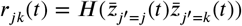, where 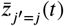 is the running average of LGN neuron *j* and 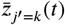 is the running average of LGN neuron *k*. The *r*_*jk*_(*t*) term ensures that only aligned LGN neurons and aligned L6 neurons are permitted to form new synapses.

Feedforward development has been studied previously (***Adelsberger-Mangan and Levy, 1993, 1994a***,b; ***Colbert et al., 1994; Thomas et al., 2015; Levy et al., 2016; Baxter and Levy, 2019, 2020***). Thus, in Results, we focus on feedback development of synapses and assume perfectly aligned feedforward connectivity from LGN to L6 exists.

### Static and retinal wave-like input patterns

The activity of the RGC neurons represents the retinal input which is restricted to a 1-D, linear retina. Retinal inputs consist of *n*_*a*_ active, adjacent RGC neurons firing for a prescribed duration, *τ*_*R*_. Each retinal input pattern is specified by a center location on the retina, *i*_*c*_, and the number of active retinal inputs, *n*_*a*_. The animal is assumed to dwell on each different retinal input pattern for a duration *τ*_*d*_.

Both binary and triangular-shaped retinal input vectors have been simulated, but in the results presented here only binary input vectors were used. When the neurons are assumed to have binary outputs and low firing thresholds, there is little difference between using binary and triangular-shaped input vectors.

Simulations with a single center location are the simplest and can be considered static inputs. Simulations that present a sequence of multiple center locations in which the active RGC neurons overlap result in more complex connectivities and weights between the LGN and L6. Retinal waves are simulated by presenting a sequence of *n*_*a*_ active RGC neurons such that their centers move from left to right and then from right to left across the field of RGC neurons. In this case, the dwell duration determines the speed of the retinal waves.

### Summary of variables and parameters

Additional parameters of the model include the number of retinal input patterns or center locations of the active retinal inputs (*n*_*c*_), and the number of training passes through all of the retinal input patterns (*n*_*p*_).

Table 1 provides a list of variables and parameters of the CTF model along with typical parameter settings.

**Table 1.**
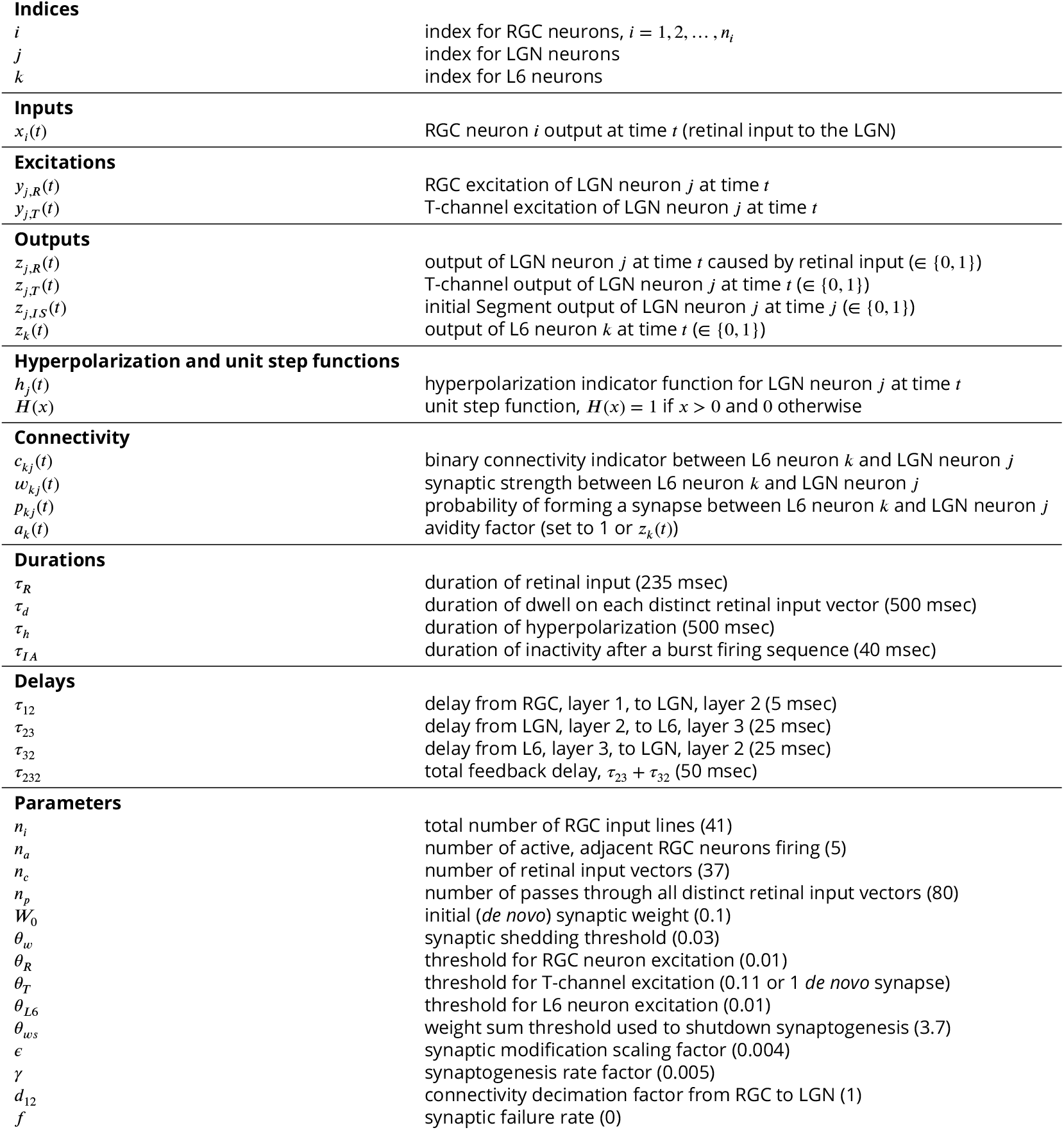
Variables and parameters. Typical parameter values are provided in parentheses.

## Results

The CorticoThalamic Feedback (CTF) model consists of three neural layers: the RGC layer, the LGN layer, and the L6 layer. The RGC represents the retinal input as a one-dimensional, linear array of equally-spaced neurons.

A small subset of the RGC neurons are active and the active RGC neurons are grouped together; i.e., the active RGC neurons are adjacent to each other. Thus, the retinal input to the LGN can be described by a center location and the number of active RGC neurons. For all of the simulations described here, the feedforward connections from the RGC to the LGN and L6 neurons are assumed to be fully developed and retinotopically aligned.

During a simulation the active RGC neurons turn on 10 msec into the simulation and turn off 235 msec later. During a simulation with active neurons located at multiple locations in the retinal array, the animal is assumed to dwell on each different retinal input for a specified duration. In the simulations presented, the dwell duration is 500 msec. After each dwell, the activities in the LGN and L6 are reset and the retinal input at a different center location is presented 10 msec into the next dwell and turned off 235 msec later. A 10 msec delay in the RGC onset and a dwell duration of 500 msec allow about nine opportunities for the signals to oscillate between LGN and L6 for each dwell.

The process of presenting the input patterns corresponding to all center locations is referred to as a pass, and multiple passes are required for the weights to reach their asymptotic values. Because the input patterns are changing, the final values of the weights may oscillate within a small range. In the simulations presented here, the center locations of the retinal input patterns progress from left to right for the first pass, from right to left for the second pass, from left to right for the third pass, and so on. The center locations shift by one RGC neuron. This sequence of retinal inputs is meant to simulate retinal waves moving across the retina.

### Spindle event timing

Figure 2 shows an example of spindle event timing as determined from the T-channel firings of the same LGN neuron. Over a short time period (as shown in Figure 2a), individual firings of the same LGN neuron are observed while the same retinal input vector is presented for the dwell duration (500 msec in this example). The temporal spacing between individual T-channel firings is determined by the LGN→L6→LGN feedback loop time (25 msec + 25 msec = 50 msec in this example). A single *spindle event* is defined as the group of spatially-specific, T-channel firings separated by the feedback loop time stimulated by the same retinal input. After all distinct retinal inputs have been dwelled upon, another spindle event occurs. In the simulations, spindle events occur 18.5 seconds apart (about 3 events per minute) and they occur at the same time on each pass, but, of course, *in vivo* spindle events occur at somewhat irregular intervals.

**Figure 2.**
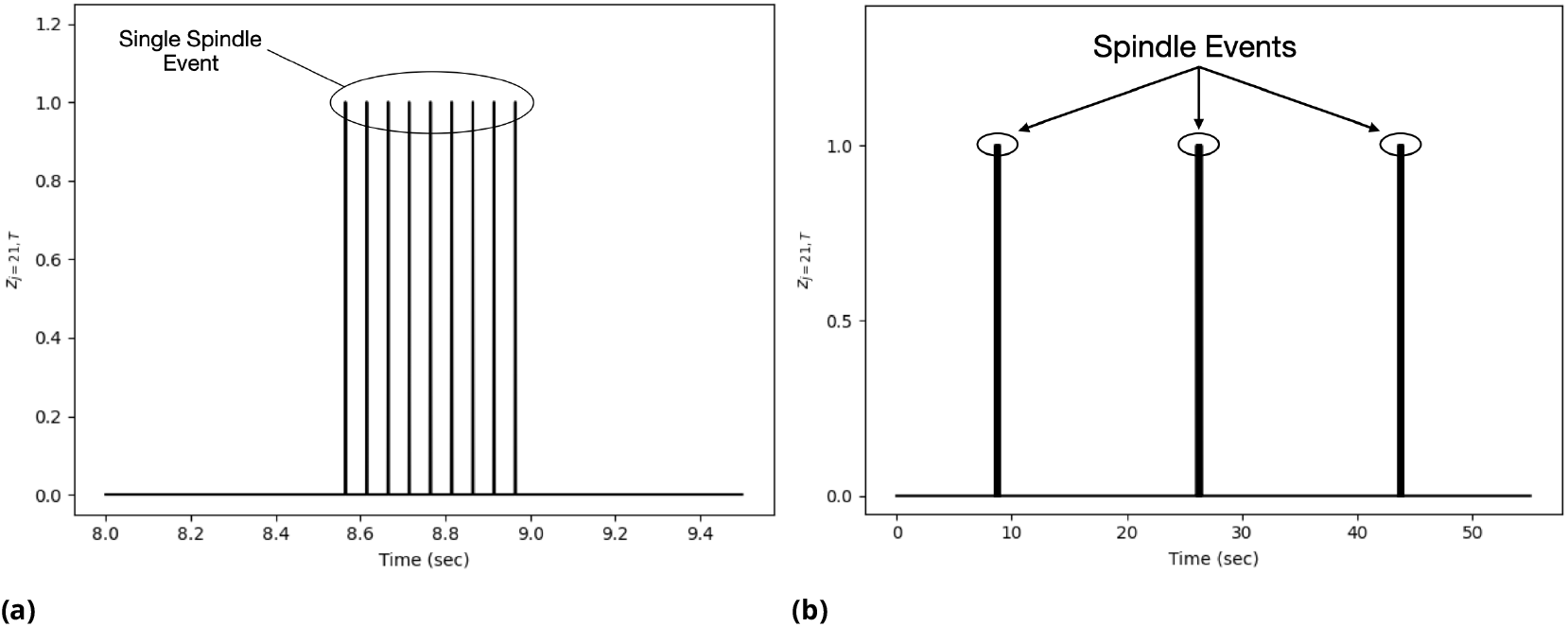
Example of simulated spindles with a spindle event consisting of nine firings of the same LGN neuron with a nominal spindle repetition of three spindle events per minute. (a) A single spindle is made up of a small number of LGN principal neurons that are being activated and then reactivated. The first firing of such neurons is due to retinal activations, and the next eight firings are due to feedback from L6 followed by T-channel activation of the same LGN neurons. Here, the LGN→L6→LGN feedback loop time is assumed to be 50 msec. Thus, the oscillation frequency of the spindle event is 20 Hz. When modeling a 25 Hz oscillation, the loop time is assumed to be 40 msec. (b) With an expanded time scale, one observes multiple spindle events from the same neuron due to the slow movement of the retinal wave that is activating LGN principal neurons. In this case, the spindle events are 18.5 seconds apart. The separation between spindle events is governed by the retinal wave dwell, a duration somewhat arbitrarily assumed to be 500 msec. (In the simulations, the size of the retina limits the number of distinct input vectors that can be simulated and dwelled upon, which, in this case, is 37 retinal positions. With this timing, one pass of the retinal wave takes 18.5 seconds and 80 retinal wave passes represents about 25 minutes of development. In biological systems, there are typically thousands of retinal wave passes and this developmental process takes three to five days.)

### Convergence of feedback weights

Given stable retinal inputs, the synaptic weights must converge to stable values to establish a stable neural code. In this section, the values of the asymptotic synaptic weights are examined in the face of changing retinal input statistics.

To examine the asymptotic synaptic weights, we start by taking the expectation of both sides of equation 8 and treating *c*_*kj*_ and *w*_*kj*_ as constants,

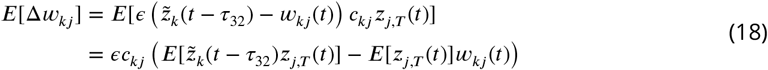

Setting *E*[Δ*w*_*kj*_ ] = 0 yields

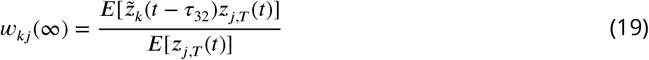

The term 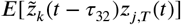 is the statistical cross correlation of 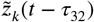 and *z*_*j,T*_ (*t*) with 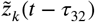 and *z*_*j,T*_ (*t*) treated as random variables. 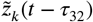 and *z*_*j,T*_ (*t*) are stationary and ergodic, the term 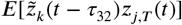 is equal to the temporal cross correlation of 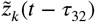 and *z*_*j,T*_ (*t*). In addition, if *E*[*z*_*j,T*_ (*t*)] > 0, the weights converge to the temporal cross correlation of the presynaptic signal, 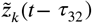, and the postsynaptic signal, *z*_*j,T*_ (*t*), normalized by the temporal average of the postsynaptic signal.

The temporal cross correlation of 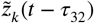 and *z*_*j,T*_ (*t*) is defined over a time interval [*t*_1_, *t*_2_] as

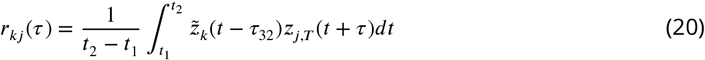

where *τ* is the time lag from 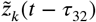 to *z*_*j,T*_ (*t*). Typically, *r*_*kj*_ (*τ*) peaks at a specific value of *τ*; the value of *τ* at the peak indicates the best temporal alignment of the two signals.

The bounding values of the asymptotic weights are determined by examining the bounds of the numerator of equation 19. If 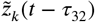 and *z*_*j,T*_ (*t*) are not appropriately timed (i.e., temporally uncorrelated), then equation 8 drives *w*_*kj*_ to zero. Alternatively, when de-inactivation and presynaptic firing always co-occur in time (i.e., perfectly correlated in time), the numerator is maximized and has a value 1. In this case, the weight is maximized and has a value of 1/*E*[*z*_*j,T*_ (*t*)]. Thus, the asymptotic value of the synaptic weight *w*_*kj*_ is bounded by the closed interval [0, 1/*µ*] where *µ* = *E*[*z*_*j,T*_ (*t*)] > 0.

Now consider the asymptotic synaptic weights when the active retinal inputs do not overlap while moving across the retina (the non-overlapping case). In this case, because the active retinal inputs do not overlap, retinal inputs corresponding to different center locations produce asymptotic values according to equation 19.

Next consider the overlapping case in which the active retinal inputs overlap while moving across the retina. In this case, there may be different asymptotic weight values at the same synapse corresponding to different center locations of the active retinal inputs. The weights in this case may oscillate within a small range of values each time the retinal input changes. From many simulations consisting of 80 passes or more, we have observed that in many cases such oscillations are negligible or do not occur; the weights appear to asymptote to values that are the average of the asymptotic weights across adjacent center locations. However, in some cases, the weights vary within a small range of values corresponding to the asymptotic weights of adjacent center locations.

Therefore, the asymptotic synaptic weights are bounded, and they are constant or stable within a small range of values in response to changes in the retinal input statistics as the retinal wave moves across the retina.

### Full model with restricted synaptogenesis

The simulation results begin with the complete model as described in the Model section. The complete and most efficient model uses restricted (spatially biased) synaptogenesis (see Equation 10). The input vectors consist of 37 center locations and each vector consists of a group of five active RGC neurons. The center locations move from left to right and from right to left on the retinal field to simulate retinal wave input. The total number of connections is graphed versus time for the restricted synaptogenesis case in Figure 3. Next, the L6→LGN weight matrix is shown at 0, 2, 5, 10, 20, and 80 passes in Figure 4. The image labeled “0 passes” is the initial connectivity, which is a mixture of appropriate (aligned) connections and random connections (for a total of 74 initial synapses). At 10 passes, the inappropriate synapses have been shed, and at 80 passes the weights in the central regions have filled in and the peaks occur along the diagonal. The profiles of the weight matrix with the L6 neuron or LGN neuron fixed at the middle of the field are graphed versus the number of passes in Figure 5. At 80 passes, the weight profiles are observed to have a triangular shape.

**Figure 3.**
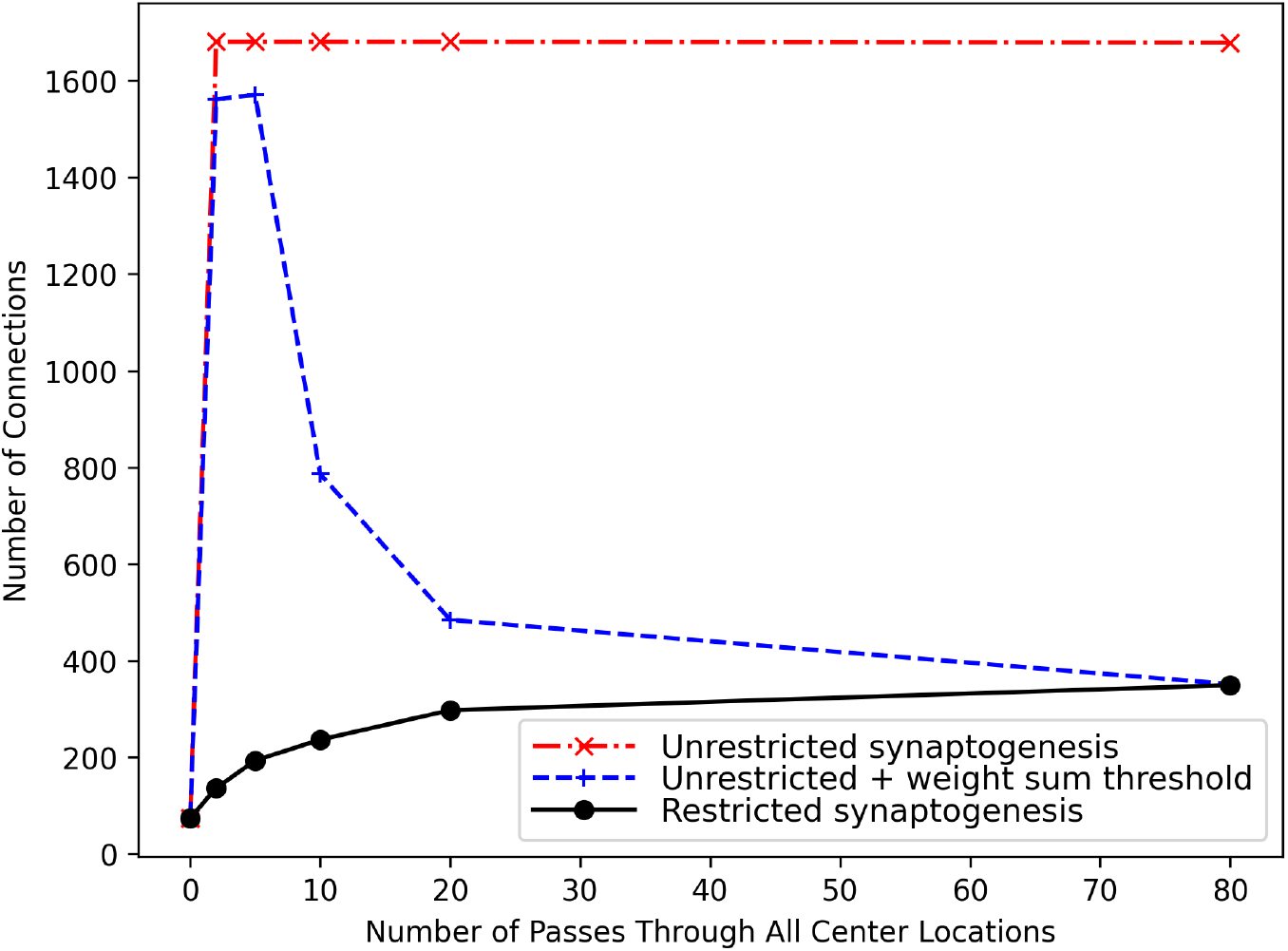
L6 to LGN connection counts versus time for the full model with biased (restricted) synaptogenesis, uniformly random (unrestricted) synaptogenesis, and uniformly random synaptogenesis with a weight sum threshold restriction (unrestricted + weight sum threshold). Equation (10), with its avidity and receptivity restrictions 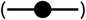, is more efficient and stable in synaptic development than (11) and (12). In fact, the completely unrestricted case (Equation 11) never stabilizes 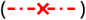. The unrestricted case with the weight sum threshold restriction (Equation 12) leads to many more de novo synapses early in development, which end up being shed 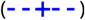. Both the restricted and unrestriced case with the weight sum threshold restriction produce nearly the same, stable number of synapses at the end. The inefficiency allowed by the unrestricted case with the weight sum threshold restriction is reflected in the initial large overshoot of synapse counts; in fact the maximum synapse count of 1571 is 93% of the maximum possible number of connections (1681).

**Figure 4.**
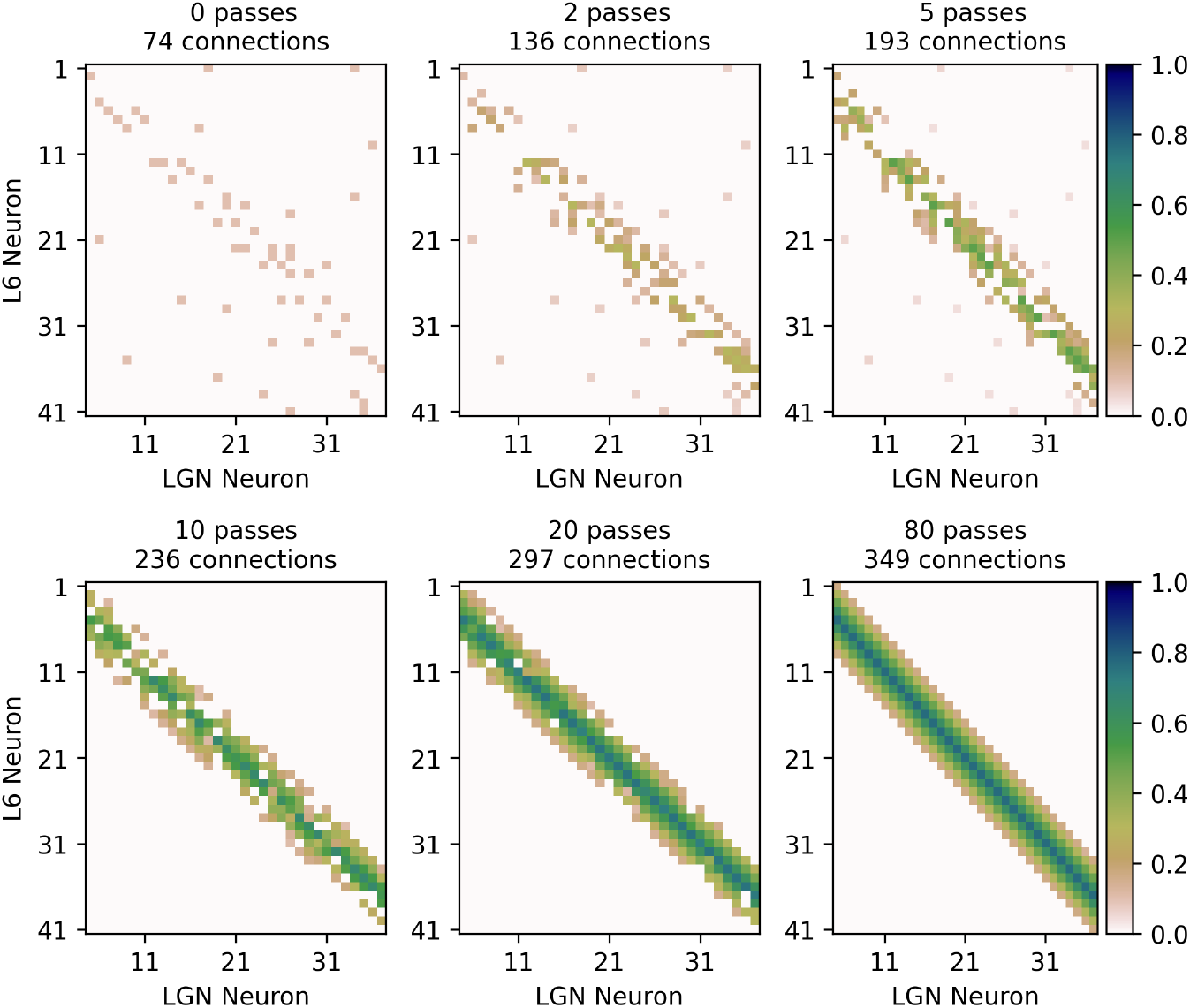
The full model, with avidity and receptivity restricting synaptogenesis (Equation 10), via feedback from V1-L6 to LGN principal neurons preserves retinotopy with mild divergence. The “0 passes” graph is the initial synaptic weights, which is a mixture of retinotopically-aligned connections and random unaligned connections. During each pass, the retinal input moved from left to right and from right to left across the retinal with five, adjacent, active RGC neurons. With successive passes, adaptive synaptogenesis increases retinotopically-aligned synaptic strengths, while unaligned connections become weaker and most are eventually discarded (shed). The retinotopic connections are found within two neurons of the diagonal of the weight matrix, i.e., where the LGN neuron index and the L6 neuron index are nearly the same. These retinotopic connections are strengthened with successive passes (brown to green). After 80 passes, synapses aligned with the active retinal input tend to get stronger; thus, synaptic weights that are aligned (near the diagonal) are larger (greener in color) compared to weights away from the diagonal that are unaligned (browner in color). Initial, randomly located, unaligned synapses (far off the diagonal) are gradually weakened and eventually shed (from brown to white). The L6 to LGN retinotopic connectivity after 80 passes is divergent with nine synapses around the diagonal rather than five synapses as would be expected with perfect retinotopy.

**Figure 5.**
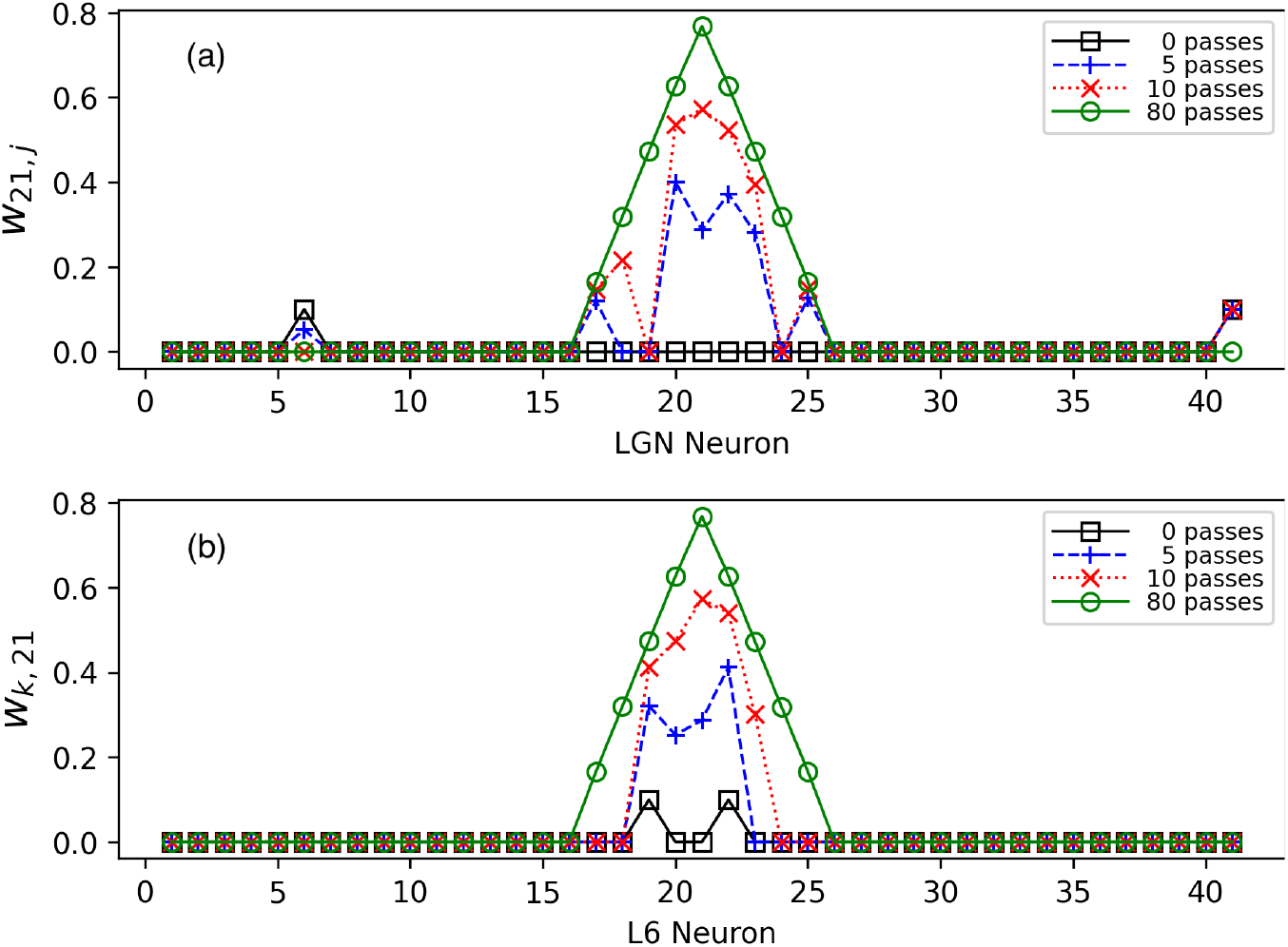
L6 to LGN synaptic development from the viewpoints of L6 neuron 21 and LGN neuron 21 (same simulation as Figure 4). The shape of the synaptic strengths is easier to visualize by graphing the synaptic weights at a fixed L6 or LGN location. (a) Synaptic weights from the perspective of L6 neuron 21. After 80 passes, the shape of the synaptic strengths is nearly symmetric and triangular. The ending number of connections is twice the width of the activity pattern minus 1; e.g., 2 × 5 − 1 = 9. Note the eventual loss of distant, unaligned synapses around LGN positions 6 and 41. (b) Synaptic weights from the perspective of LGN neuron 21. Again, after 80 passes, the shape of synaptic weights is nearly symmetric and triangular, and the number of connections is 9.

### Full model with unrestricted synaptogenesis

As Figure 3 indicates, unrestricted (random) synaptogenesis does not converge to a stable number of connections, but adding a weight sum threshold restriction provides stable connectivity. Figures 6 and 7 show the weight matrix and weight profile of an example simulation with unrestricted synaptogenesis with a weight sum threshold restriction added (see Equation 12 in Model). The weight sum restriction only requires a single parameter, the weight sum threshold, *θ*_*ws*_, and an appropriate value of this parameter can be easily estimated. With five active RGC neurons, the value that turned out to be effective was approximately 3.7, which is just below five times the asymptotic weight value of about 0.77.

**Figure 6.**
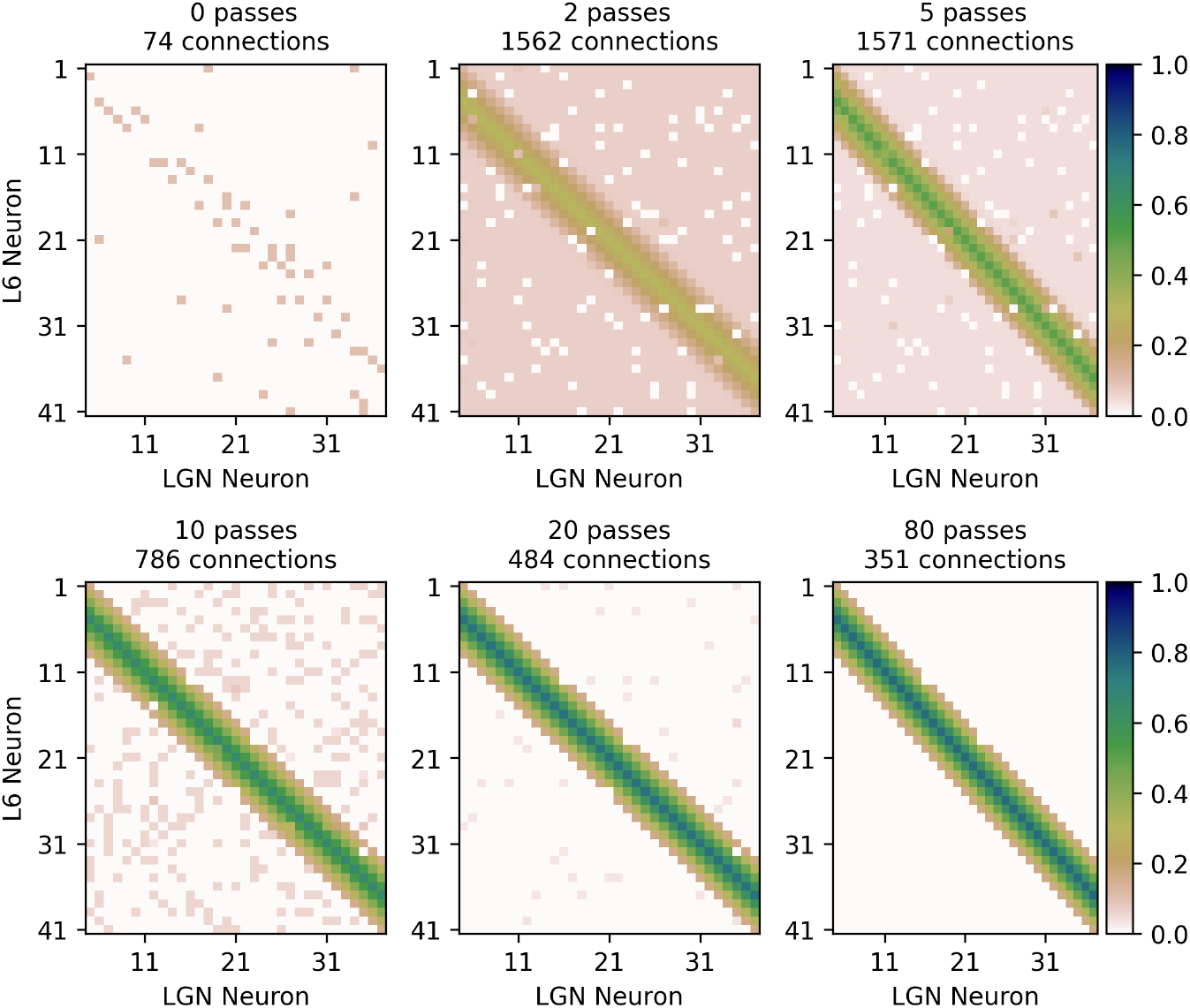
The model without avidity and receptivity restricting synaptogenesis but with a weight sum restriction (Equation 12), becomes almost fully connected early on but then evolves to preserve retinotopy with a weight matrix quite similar to that of the full model shown in Figure 4. Comparing this figure to Figure 4 at passes 2, 5, and 10 illustrates the inefficiency produced by removing the avidity and receptivity restrictions on synaptogenesis. The “0 passes” graph is the same as the one found in Figure 4, showing the initial connectivity, which is a mixture of retinotopically-aligned connections and random, unaligned connections. The pervasive light brown background found at passes 2 and 5 corresponds to the large number of synapses illustrated in Figure 3 when synaptogenesis is unrestricted except for a weight sum threshold restriction. With successive passes and given enough developmental experience, the retinotopic synapses strengthen (brown to green) while unaligned connections become weaker and most are eventually shed (brown to white). As in Figure 4, retinotopic connections are found within two neurons of the diagonal. After 5 passes, there are 1571 synapses (93% of all possible synapses), but after 80 passes most inappropriate synapses have been shed, aligned synapses have strengthened, and the peak synaptic weights are found along the diagonal. As in Figure 4, the L6 to LGN retinotopic connectivity after 80 passes has nine synapses around the diagonal rather than the five that would be expected with perfect retinotopy.

**Figure 7.**
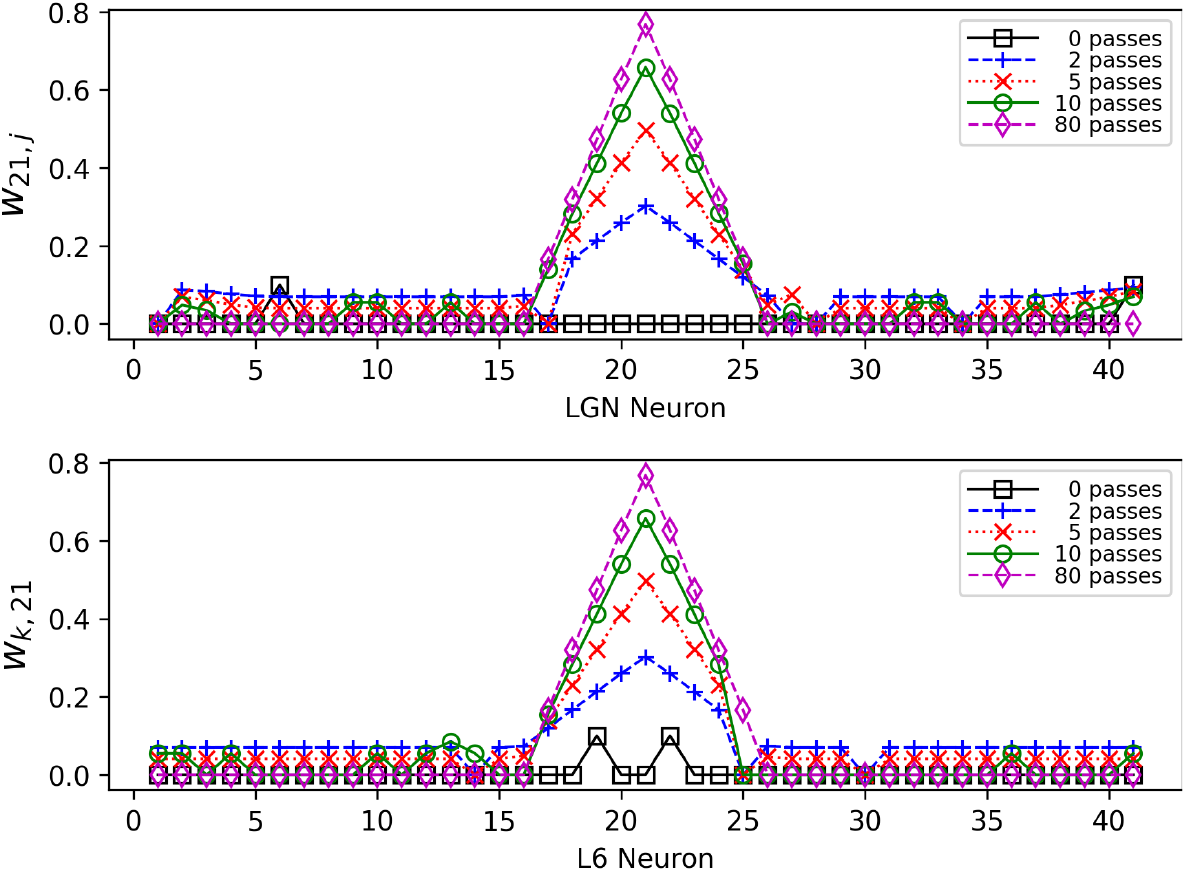
Synaptic weight profiles versus time for the model with unrestricted synaptogenesis but with a weight sum threshold restriction (same simulation as Figure 6). (a) Weight profile from the viewpoint of L6 neuron 21. After 80 passes, the shape of the synaptic strengths is nearly symmetric and triangular, and the number of connections is 9. (b) Weight profile from the viewpoint of LGN neuron 21. Again, after 80 passes, the shape of the synaptic strengths is nearly symmetric and triangular, and the number of connections is 9.

Comparing the solid, black curve in Figure 3 with the dashed, blue curve reveals that unrestricted synaptogenesis produces a significant overshoot of synapses – a maximum of 1571 synapses after five passes compared to 351 synapses after 80 passes (an overshoot ratio of about 4.5).

Figure 6 shows that at 2 and 5 passes, there are an abundance of synapses created (maximum possible is 41 × 41 = 1681). Figure 7 shows the weight profiles corresponding to the same simulation in Figure 6. As in Figure 5, Figure 7 shows that after 80 passes the weight profiles are symmetrical with respect to index 21.

Based on the differences between restricted and unrestricted synaptogenesis, *restricted synaptogenesis is observed to be the most efficient method of developing connectivity with the least amount of unnecessary connections and more steady and stable connectivity during development*.

### Varying the number of active RGC neurons

Figure 8 shows the effect of the number of active RGC neurons on the synaptic weights and synapse counts. In Figure 8a, the synaptic weights are shown with the L6 index fixed at 21. The spread of the synaptic weights increases with an increasing number of RGC neurons. Figure 8b shows that the number of synaptic connections is an approximately linear function of the number of active RGC neurons.

**Figure 8.**
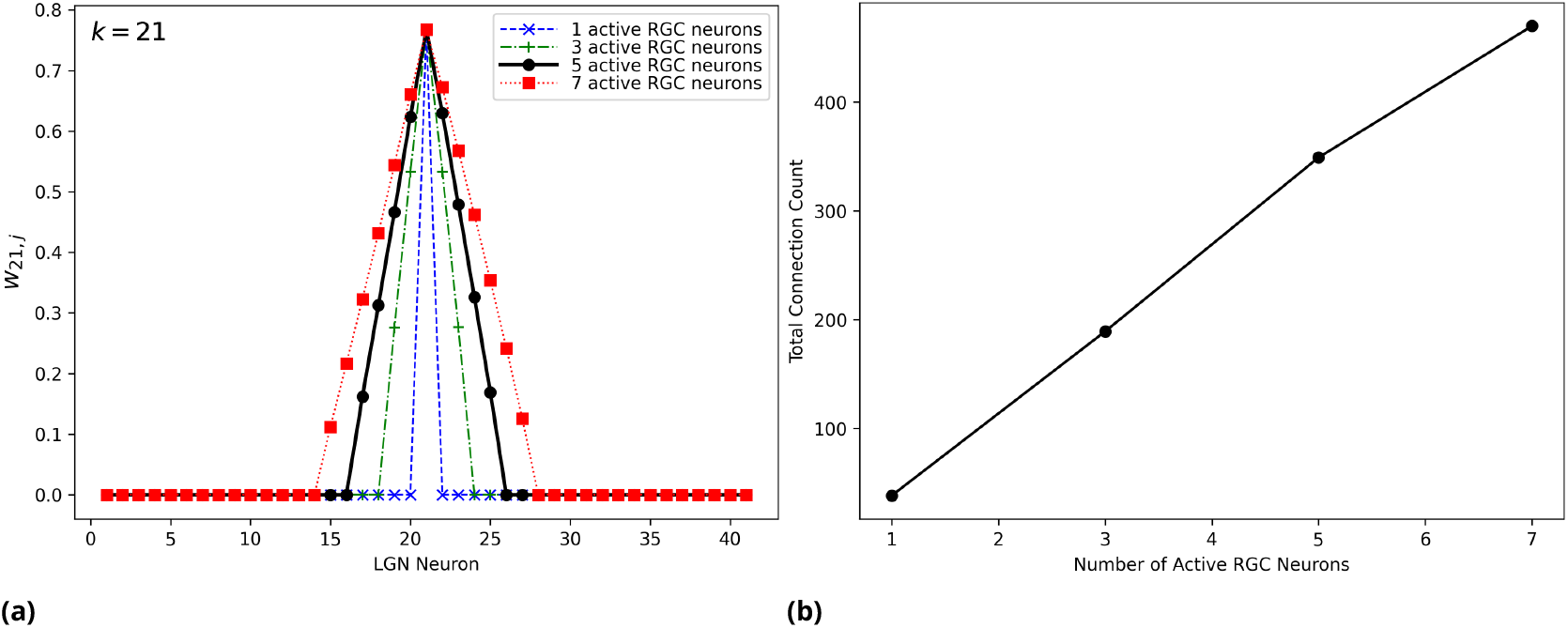
Developed synaptic weights and synapse counts versus the number of active RGC neurons. (a) Synaptic weights from the perspective of L6 neuron 21 after asymptotically developing (80 passes). Feedback retinotopy is divergent, and divergence increases with the number of active RGC neurons. The synapse counts culminate in one less than twice the number of active RGC neurons; e.g., if retinal activity consists of 5 RGC neurons, the count is 9 (2 × 5 − 1). (b) The total synapse counts versus the number of active RGC neurons. The number of connections increases approximately linearly with the increasing number of RGC neurons. These simulations are based on the full model with hyperpolarization and the T-channel with simulated retinal wave input (37 distinct retinal input vectors moving across a retina consisting of 41 RGC neurons).

### Varying the number of active L6 neurons

Figure 8 varied the number of active input lines (RGC neurons) in the input vectors. In contrast, one can selectively inhibit all but *K* L6 neurons and observe the resulting development of weights. Such inhibition could be the result of lateral inhibition or the selection of *K* L6 neurons with the highest excitations. With *K* = 1, all L6 neurons were inhibited except for the center L6 neuron; with *K* = 3, the three L6 neurons in the center were permitted to fire and the remaining neurons were inhibited; with *K* = 5, the five L6 neurons were permitted to fire and the remaining neurons were inhibited.

Figure 9 shows the weight profile and connection counts with a varying number of active L6 neurons using this L6 selective inhibition procedure. In Figure 9a, the solid, black curve corresponds to the case without L6 selective inhibition; in this case, the number of active L6 neurons equals the number of active RGC neurons. The other two curves correspond to the inhibition of two or four L6 neurons which produces three or one active L6 neuron. Fewer active L6 neurons lead to significantly diminished peak synaptic strengths. Figure 9b shows that the number of connections is a linear function of the number of active L6 neurons.

**Figure 9.**
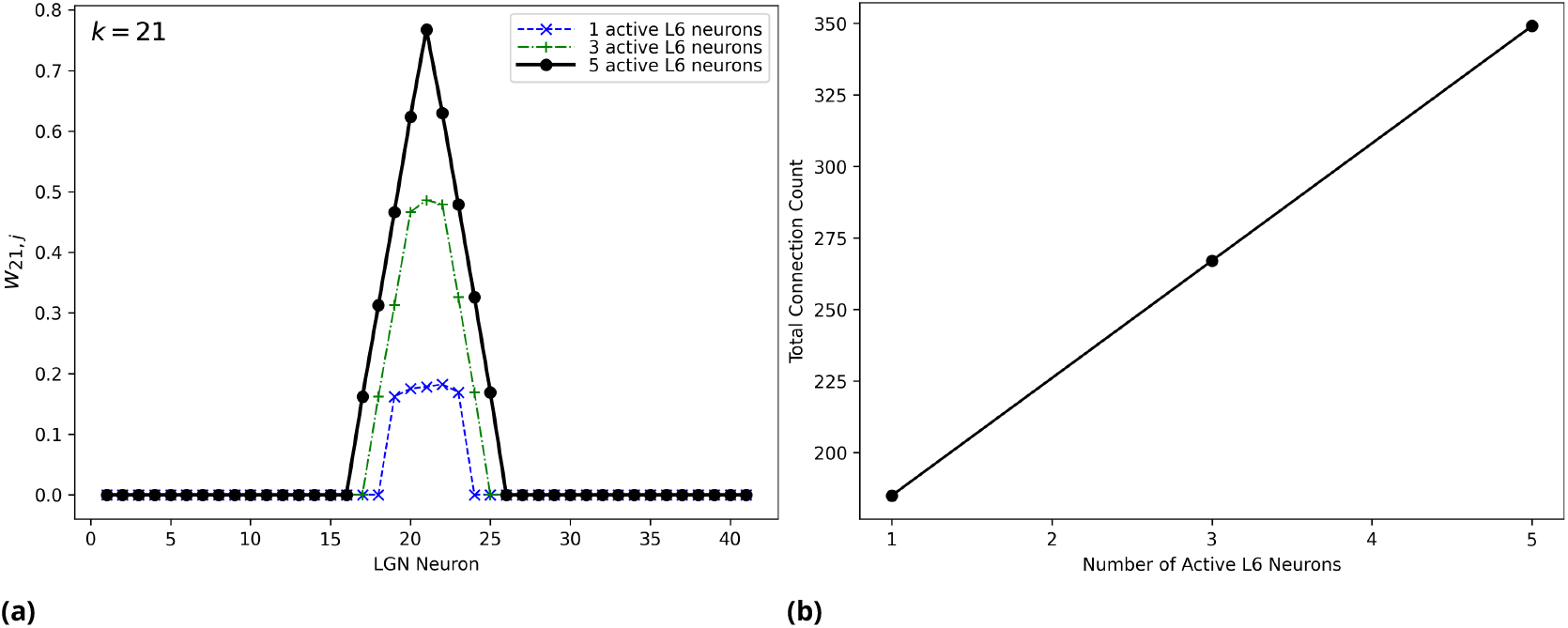
Developed synaptic weights and synapse counts versus the number of active L6 neurons. (a) Developed synaptic weights with the L6 neuron index fixed at 21 versus the number of active L6 neurons. Fewer active L6 neurons significantly diminish the peak synaptic strengths. Here, the 5 active L6 neurons case 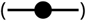 is the same as the case without L6 selective inhibition found in Figure 8, i.e., the number of active L6 neurons is equal to the number of active RGC neurons. The 3 active L6 neurons 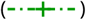 case and 1 active L6 neuron 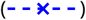 case are both impacted by inhibition, i.e., two and four of the L6 neurons are inhibited, respectively. (b) The synapse counts versus the number of active L6 neurons is approximately linear. These simulations are based on the full model with hyperpolarization and the T-channel with simulated retinal wave input (37 distinct retinal input vectors moving across a retina consisting of 41 RGC neurons).

### Varying output functions and firing probabilities

Three types of output functions were simulated: binary with a specified firing threshold, threshold-linear with a specified threshold for the transition to linear output, and sigmoid with the steepness and the input value that produces an output of 0.5 specified. Because a threshold-linear function is not bounded, it tended to produce unstable connectivity and was deemed inappropriate for the feedback loop. With appropriate parameter adjustments, the sigmoid function produced similar results to the binary function.

We have also found that varying the firing probabilities of the active LGN and L6 neurons changes the spread of the weight profiles.

### Receptivity and avidity have a strong effect on synaptogenesis

Recall that synaptogenesis is the product of avidity, receptivity, and the synapse formation rate parameter *γ*. Avidity is associated with the presynaptic signal and receptivity is associated with the postsynaptic signal. Figure 10 shows how the connectivity develops for the full model with restricted synaptogenesis, with restricted synaptogenesis but no avidity, with restricted synaptogenesis but no receptivity, and with unrestricted synaptogenesis and a weight sum threshold restriction but no avidity. Here, “no avidity” is an abbreviated way of indicating that the avidity function is set to 1 so that synaptogenesis is independent of avidity. The term “no receptivity” means that receptivity is set to 0, so that synaptogenesis is turned off for the entire simulation; in this case, no new connections are permitted and the connectivity is determined by the initial connectivity and shedding. The initial connectivity can be thought of as the chemotactic connectivity.

**Figure 10.**
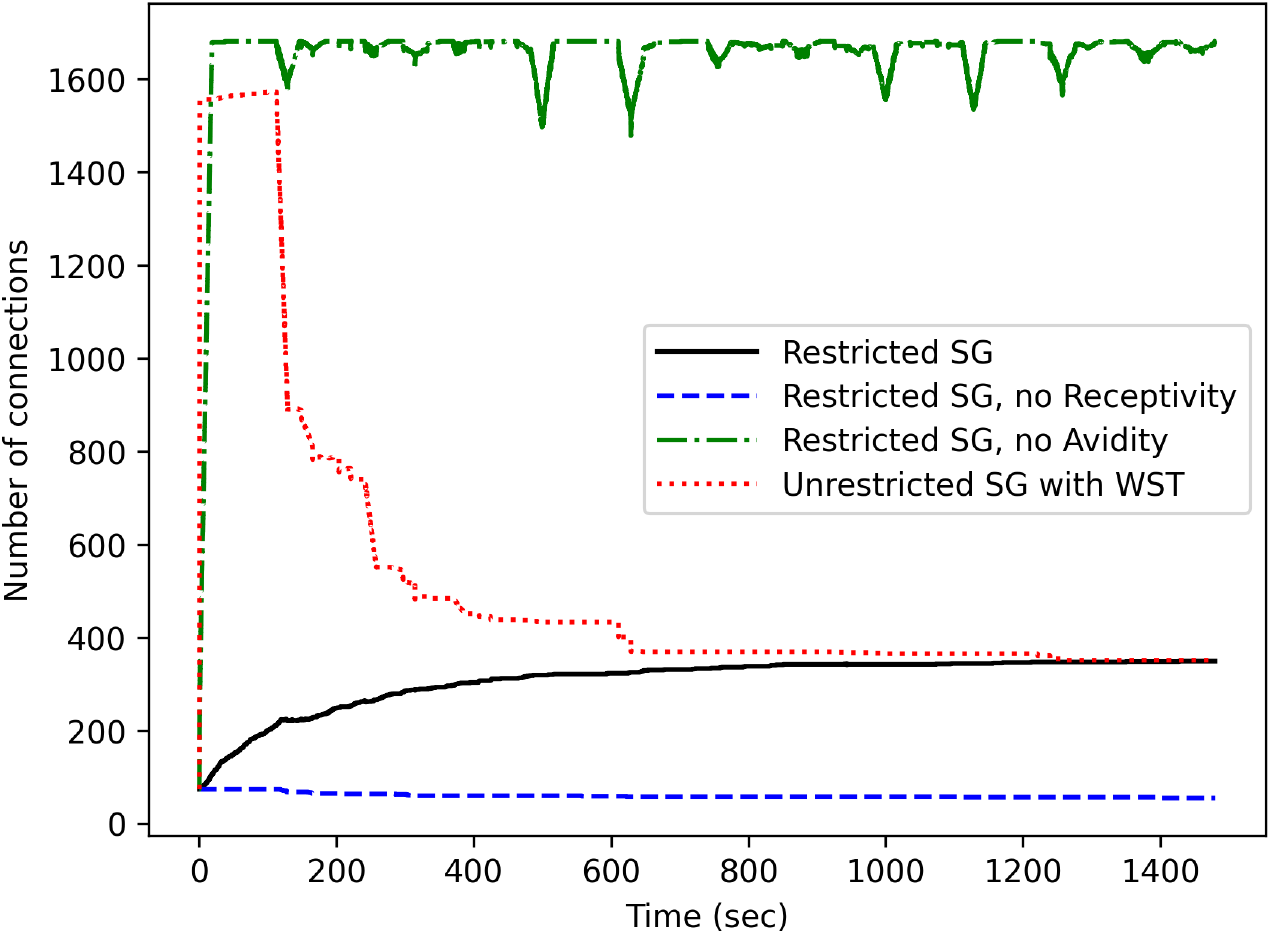
L6 to LGN connection counts versus time for the adaptive synaptogenesis model with four different types of synaptogenesis restrictions. Restricted synaptogenesis with avidity and receptivity 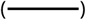 has the most stable and efficient synaptic development. Restricted synaptogenesis without avidity 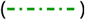, i.e., with avidity set to 1, produces unstable connections with nearly 100% connectivity. In the restricted synaptogenesis case without receptivity 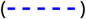, i.e., with receptivity set to 0, there are no new connections, although there are a few shedding events. Unrestricted synaptogenesis case with a weight sum threshold restriction 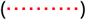 produces a large initial overshoot, given avidity is set to 1, but the weight sum threshold restriction stabilizes the connectivity such that unnecessary connections are eventually shed, producing nearly the same number of synapses at the end as the restricted case. SG = synaptogenesis, WST = weight sum threshold restriction.

The solid, black curve in Figure 10 shows the full model with restricted synaptogenesis, which is similar in shape to the solid, black curve in Figure 3. The only difference in these curves is the x-axis: one is graphed as a function of time and the other is graphed as a function of the number of passes. These curves show very little, if any, overshoot of synapse counts which implies efficient development of connectivity.

The dashed-dotted, green curve shows the full model with restricted synaptogenesis with avidity set to 1, which is similar in shape to the dashed-dotted, red curve in Figure 3 which represents unrestricted (uniformly random) synaptogenesis. Thus, we can see that setting avidity to 1 versus setting avidity to *z*_*k*_(*t*) has a dramatic effect on the stability of connections. As we will see, the unstable connections are the unaligned connections.

Now compare the dotted, red curve in Figure 10 to the dashed, blue curve in Figure 3 (note the difference in the x-axes). Both of these two curves represent the full model with unrestricted synaptogenesis with avidity set to 1 and with the weight sum threshold restriction. Note that the ending synapse counts are nearly identical to restricted synaptogenesis with avidity and receptivity (the solid, black curve).

And finally observe the dashed, blue curve in Figure 10 which represents restricted synaptogenesis with receptivity set to 0. In this case, new connections are not permitted but some connections are shed. The simulation starts out with 74 initial connections, sheds 19 unaligned connections, and ends up with 55 connections. Compared to the full model with avidity and receptivity that ends up with 349 connections, the connectivity in this case is quite inadequate.

### Synaptogenesis with a spatial distance restriction

In Model, six methods of stopping synaptogenesis were described. We used the method based on limiting the sum of the weights to each LGN neuron (equation 12) in Figures 6 and 7, and in the blue, dashed curve in Figure 3. While this mechanism is effective, it is often not necessary with restricted synaptogenesis because after several passes through all of the input vectors located at each center the appropriate number of synapses is obtained and synaptogenesis turns off without any additional shutoff mechanism.

Now let’s examine the effect of adding a spatial distance restriction to unrestricted synaptogenesis. Figure 11 graphs generalized Gaussian spatial distance attenuation functions with a distance threshold of 6 for *β* parameters 1, 1.5, 2, and 4. The distance attenuation functions are designed to have a non-zero minimum attenuation, denoted by *d*_min_, so that connections between distant neurons are permitted with a low but non-zero probability. The distance threshold, denoted by *θ*_*d*_, determines the distance at which the minimum attenuation applies. Neurons with a spatial separation less than *θ*_*d*_ have a moderate to high probability of synapse formation.

Figure 12 shows the connection counts versus time for unrestricted synaptogenesis with spatial distance restrictions using distance thresholds of 3, 5, and 10. In these simulations, the generalized Gaussian distance attenuation function was used (equation 14) with *d*_min_ = 10^−6^ and *β* = 1.5. A spatial distance threshold of 5 produces connectivity and weights almost identical to the restricted synaptogenesis case except that a modest overshoot of synapses is observed compared to the case without the distance restriction.

**Figure 11.**
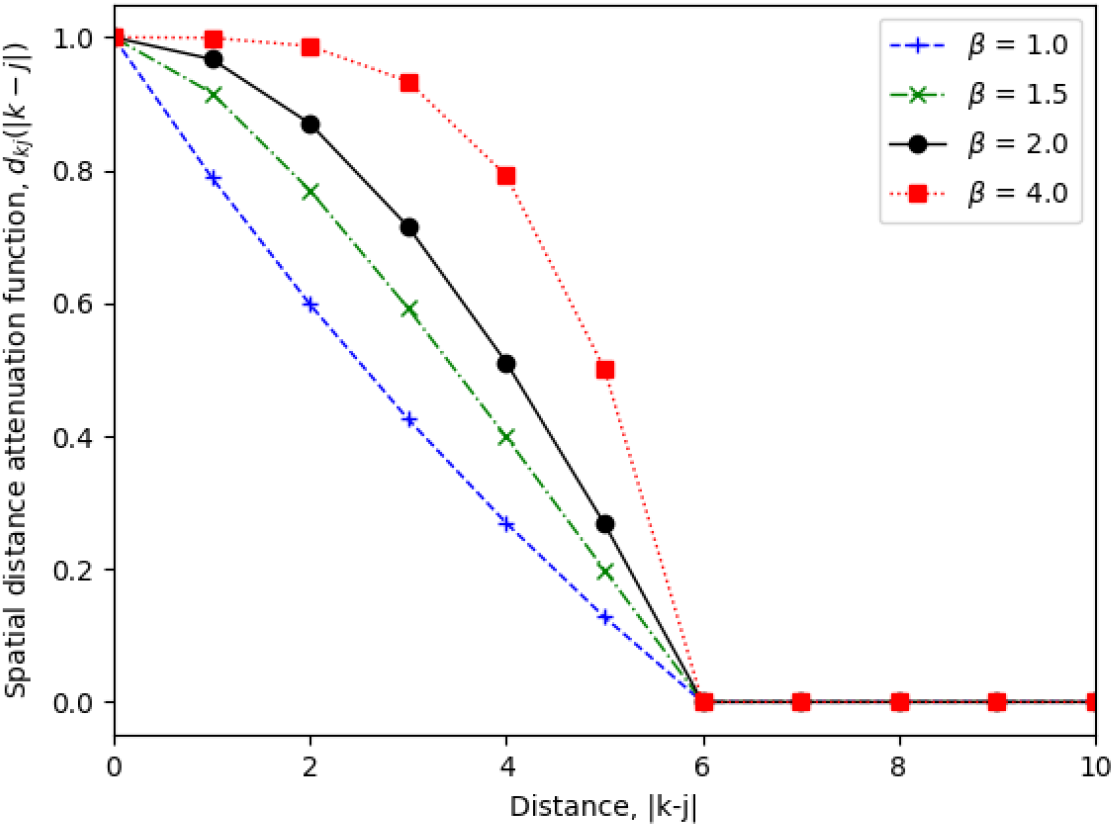
The spatial distance attenuation function *d*_*kj*_ (|*k* − *j*|) (equation 14) attenuates the synapse formation probability (see equation 13) based on the distance between the L6 neuron *k* and LGN neuron *j*. The distance is assumed to be related to the absolute difference between the neuron indices, |*k* − *j*|. Here, generalized Gaussian distance attenuation functions are graphed as a function of the distance |*k* − *j*| for several values of the shape parameter *β* (*β* = 2 is a Gaussian function and *β* = 1 is a Laplacian). With *β* ≤ 1 the function is concave upward; with *β* > 1 the function is concave downward. In Figures 12, 13, and 14 *β* = 1.5. The distance threshold, *θ*_*d*_, determines when the distance function decreases to the minimum attenuation *d*_min_; i.e., *d*_*kj*_ = *d*_min_ when |*k* − *j*| ≥ *θ*_*d*_. In this figure, *θ*_*d*_ = 6 and *d*_min_ = 10^−6^.

**Figure 12.**
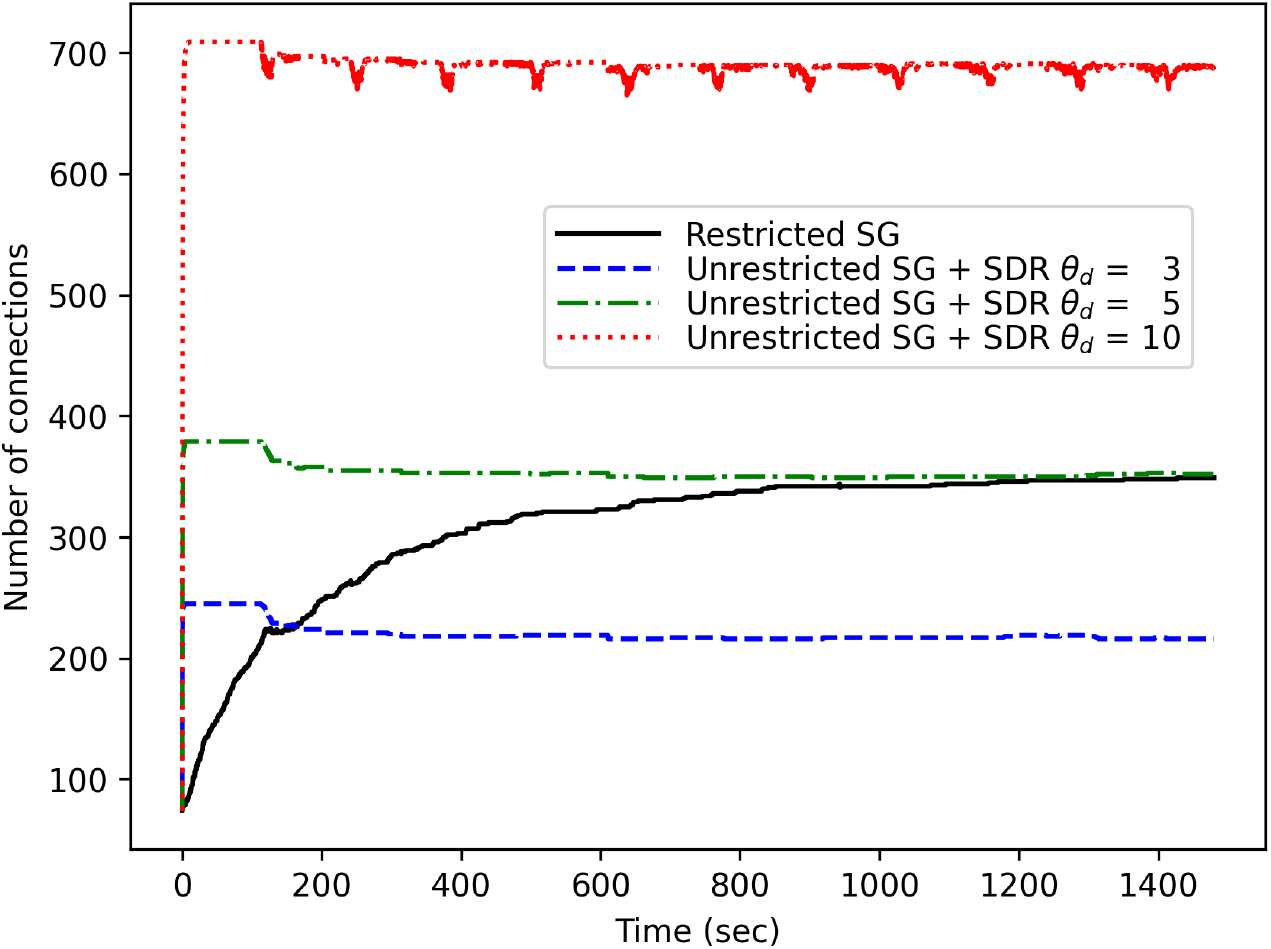
Connection counts versus time for unrestricted synaptogenesis with different spatial distance restrictions using distance thresholds *θ*_*d*_ = 3, 5, and 10. The spatial distance attenuation function with *β* = 1.5 and *d*_min_ = 10^−6^ is used (see Figure 11). In the unrestricted synaptogenesis cases, as *θ*_*d*_ increases the number of connections increases, i.e., the case with *θ*_*d*_ set to 10 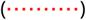 has the greatest number of connections, while the case with *θ*_*d*_ set to 3 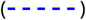 has the least. The curve representing unrestricted synaptogenesis with *θ*_*d*_ set to 5 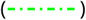 produces approximately the same number of connections as the restricted synaptogenesis case 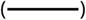, however, in the unrestricted case, there is a modest overshoot of synapses and slight shedding. The number of active RGC neurons is 5. SG = synaptogenesis, SDR = spatial distance restriction.

Figure 13 compares the ending weight matrices for restricted synaptogenesis and those for three spatial distance thresholds. The ending weights for the restricted synaptogenesis case are almost identical to those with a spatial distance threshold of 5 (the retinal inputs had five active RGC neurons). With a distance threshold of 1 (not shown), the connectivity is mostly confined to the diagonal; with a distance threshold of 10, the connectivity spreads beyond five synapses into the unaligned region which causes unstable connectivity (an endless cycle of the creation of new synapses in the unaligned region followed by shedding). With a spatial distance threshold of 3, the ending synaptic weights are located mostly within two neurons from the diagonal (five non-zero weights near the diagonal), which is the same number of connections as the number of active RGC neurons.

**Figure 13.**
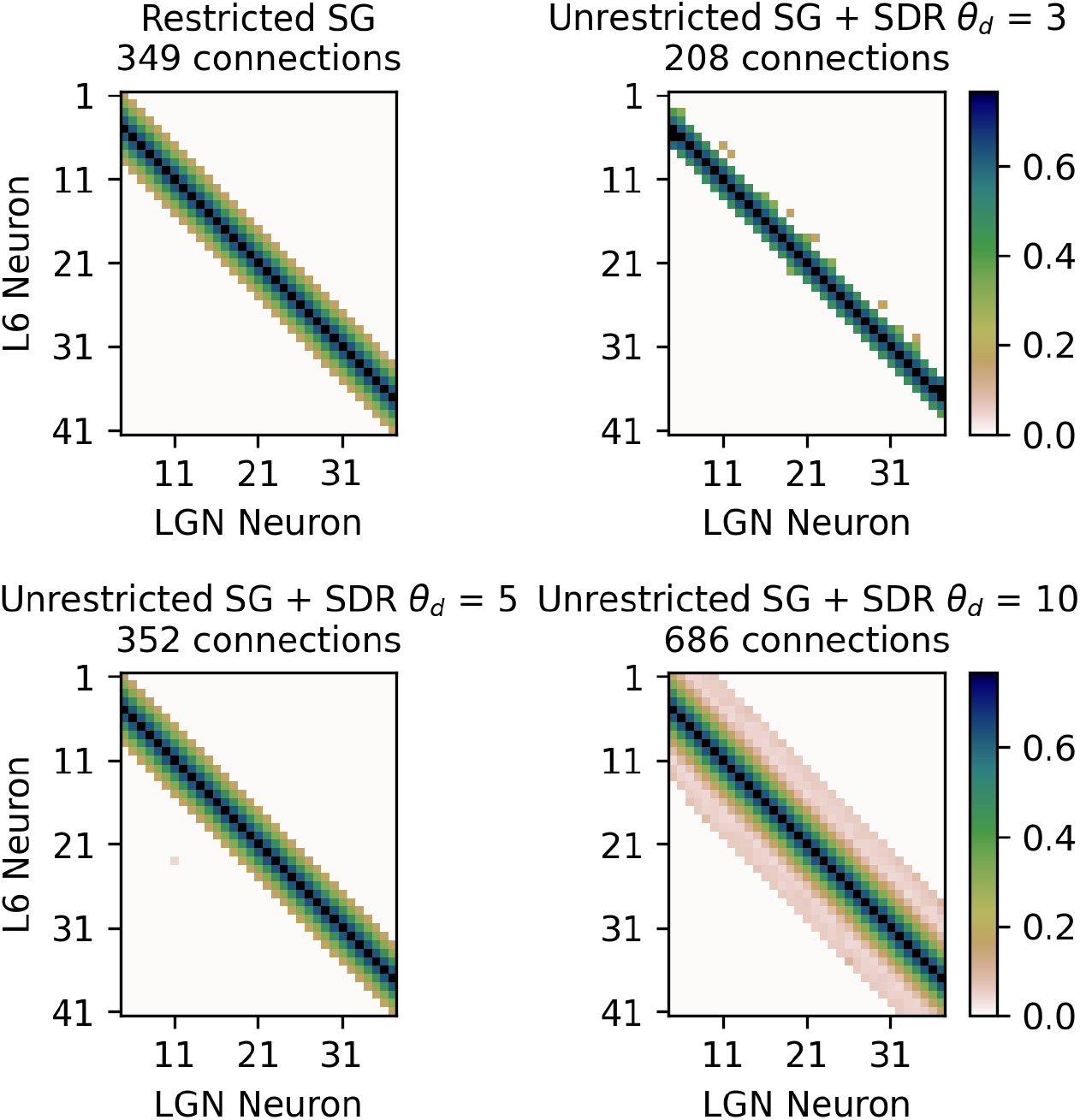
Comparison of the ending weight matrices for unrestricted synaptogenesis with different spatial distance thresholds (*θ*_*d*_) and the ending weight matrix for restricted synaptogenesis. The ending weights for the unrestricted synaptogenesis case with *θ*_*d*_ = 5 is nearly the same as the restricted synaptogenesis case, i.e. the retinotopic connectivity is divergent with nine synapses around the diagonal and the weight values are nearly identical. For example, in both cases, activation of position 21 will activate position 21 and co-activate neighboring positions 17 through 25 (± 4). The larger the distance threshold, the more divergent the connectivity, i.e., it is less aligned (further from the diagonal). The case of unrestricted synaptogenesis with *θ*_*d*_ of 3 produces 5 connections within two neurons from the diagonal, which is the same number of connections as the number of active RGC neurons. The unrestricted synaptogenesis case with *θ*_*d*_ of 10 produces highly divergent connectivity, and the connections outside the aligned region are unstable. SG = synaptogenesis, SDR = spatial distance restriction.

Figure 14 provides the weight matrices at 0, 2, 5, 10, 20, and 80 passes with a distance threshold of 3. There is very little difference between the weights at 10, 20, and 80 passes. The connections are mostly along the diagonal or within two neurons of the diagonal, and the strongest weights are all along the diagonal. The weakest weights in the aligned region are two neurons from the diagonal and are 60% of the strength of the weights along the diagonal.

**Figure 14.**
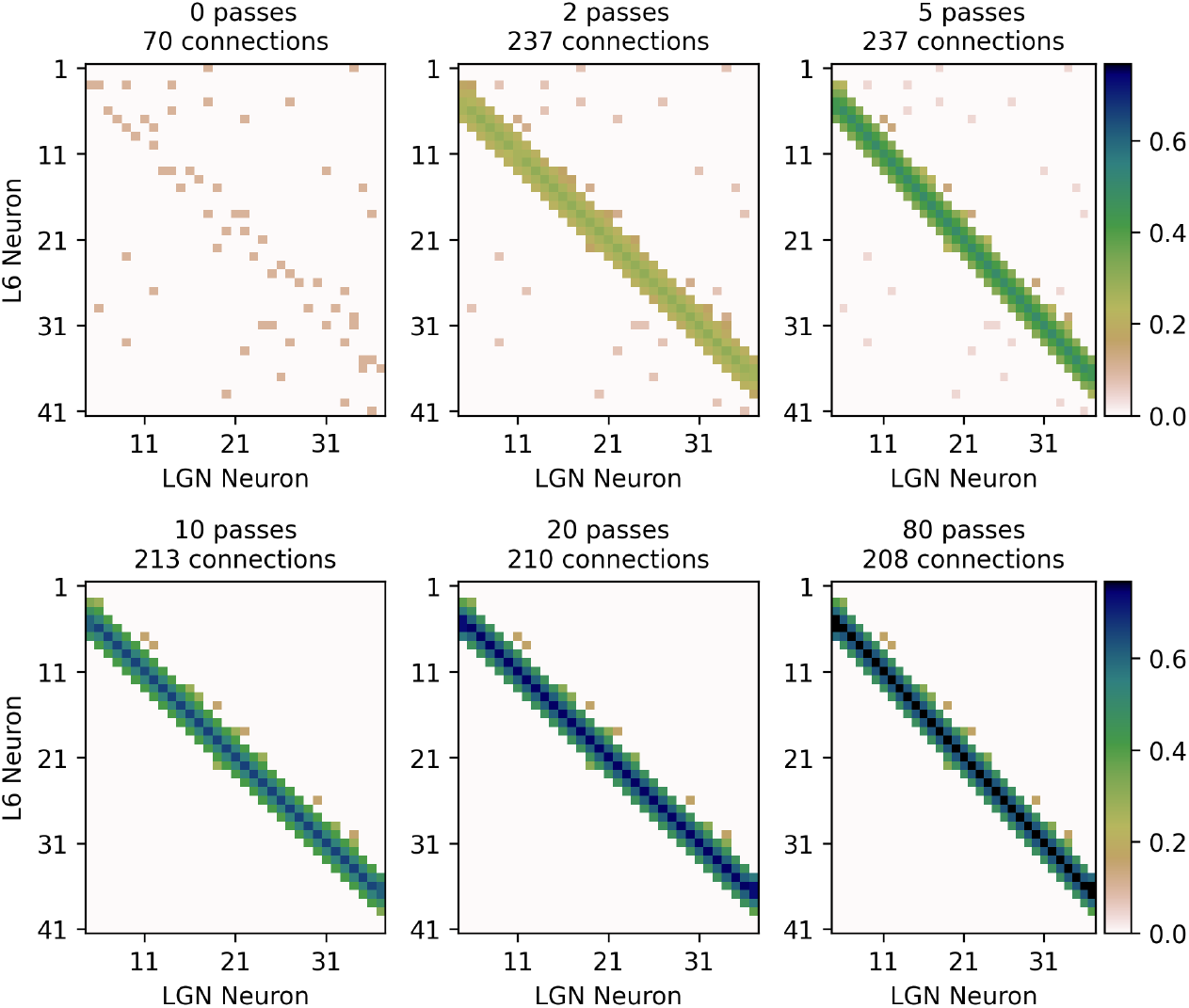
The full model with unrestricted synaptogenesis but with a spatial distance restriction (see equation 13) with a distance threshold of 3 (*θ*_*d*_ = 3) evolves to develop feedback from V1-L6 to LGN principal neurons with nearly perfect retinotopy. Here, perfect retinotopic connectivity consists of connections within two neurons of the diagonal because there are 5 active RGC neurons. In this case, there are a few unaligned weak connections after 80 passes. The synaptic weights two neurons from the diagonal are approximately 60% of the strength of the diagonal weights. The “0 passes” graph is the same as the one found in Figures 4 and 6 showing the initial connectivity, which is a mixture of retinotopically-aligned connections and random, unaligned connections. With successive passes, adaptive synaptogenesis increases retinotopic connectivity while the highly divergent connections become weaker and are eventually discarded (shed). After 10 passes, most of the connections are retinotopic. After 80 passes, synaptic weights that are aligned (within two neurons of the diagonal) are larger (darker in color), with the largest weights on the diagonal, compared to weights more than two neurons away from the diagonal that are unaligned (mostly brown in color). There is very little difference between the weight matrices at 10, 20, and 80 passes. The initial, randomly located, unaligned synapses (far off the diagonal) are gradually weakened and most are eventually shed (from brown to white). (Simulated retinal wave input with 37 distinct retinal input vectors moving across a retina consisting of 41 RGC neurons).

## Discussion

### Summary of results

We have described a minimal model that uses synaptogenesis, Hebbian/anti-Hebbian modification of existing synapses, and synaptic shedding to establish retinotopic feedback connections from layer 6 of the primary visual cortex (area V1) to the LGN. These L6→LGN connections are approximately reciprocal to the feedforward LGN→L6 connections; i.e., |*w*_*kj*_ − *w*_*jk*_| ≤ *δ* where *δ* is a small, nonnegative value. Once these reciprocal connections have been established, they are maintained in the face of changing retinotopic activity and changing cortical activity.

The model takes into account the delayed signals inherent in the feedback loop. We have shown that essentially the same synaptic weight update rule can be used for feedforward synapses and feedback synapses. However, while the feedforward synaptic modification rule uses the NMDA receptor, the feedback rule uses hyperpolarization of LGN principal cells and T-channels. In order to preserve retinotopy, there must exist a memory of the specific retinotopic excitation. This memory of the recent retinotopic excitation is provided by the time course of hyperpolarization, and subsequent T-channel bursting in response to synaptic activation. Such a sequence has been called rebound firing and exists in more than one form (***Grenier et al., 1998***).

The restricted synaptogenesis rule (equation 10) allows new synapses to form only when the T-channel is not firing, while the synaptic modification rule (equation 8) requires T-channel activation and, thus, one or more LGN firings. Figure 4 demonstrates how these two rules work together to efficiently form appropriate connections and strengthen appropriate synapses. Figure 3 demonstrates that the restricted synaptogenesis rule produces extremely efficient synapse formation with a minimal amount of unneeded connections. Figure 3 also demonstrates that with the unrestricted synaptogenesis rule (equation 11), additional restrictions are required to achieve stable connectivity.

Figure 8 shows how increasing the spatial extent of retinotopic activity increases the spatial (retinotopic) spread of synaptic weights. Thus, the number of feedback connections increases in a nearly linear fashion as a function of retinotopic activity. Figure 9 assumes that there is some sort of cortical lateral inhibition that inhibits the retinotopic extent of L6 activity leading to smaller synaptic weights, fewer connections, and a decreased spread of the weights.

The simulations show that there is a tendency for some small amount of retinotopic divergence, which may or may not be physiological. That is, as far as we know, there is no anatomical data in literature measuring the retinotopic precision of feedback for the L6 to LGN system.

A seemingly necessary part of our hypothesis is a biophysical compartmentalization such that excitation of retinal synapses on LGN neurons cannot produce T-channel mediated burst firing (***Mukherjee and Kaplan, 1995***). Rather it is only the dendritically localized excitatory, feedback synapses that have access to the burst-firing, T-channel mechanism.

### Curtailing synaptogenesis

The receptivity term in the synaptogenesis equation (equation 10) helps to curtail synaptogenesis. Synaptogenesis is limited to those LGN neurons that are not sufficiently innervated. When the postsynaptic, i.e. LGN neuron, is burst firing there is no reason for further synaptogenesis. Figure 10 shows that without avidity the connectivity is unstable so avidity also plays a role in limiting synaptogenesis.

When the model is run with unrestricted (uniformly random) synaptogenesis, connections continue to form and are shed throughout the simulations (see the red, dashed-dotted curve in Figure 3). To establish stable connectivity with unrestricted synaptogenesis, additional restrictions are required to stop synaptogenesis. In Model, five additional mechanisms are listed to curtail synaptogenesis for the unrestricted case. Two of these five additional mechanisms for shutting down synaptogenesis for the unrestricted model were demonstrated in Results: (a) a threshold on the sum of the weights and (b) a spatial distance restriction.

Biologically, the sum of the weights is related to the number of AMPA-receptors so a threshold on the sum of the weights is certainly local and plausible. The spatial distance restriction results in a higher synapse formation probability when neurons are close and a lower synapse formation probability when neurons are far away, which is consistent with biological observations.

In the biological context, eye-opening is correlated with less spindling (***Shen and Colonnese, 2016; Rochefort et al., 2009***). If we suppose that synaptogenesis is spindle-dependent then this lessening of spindle events would lessen the amount of synaptogenesis.

### Related research

***Butts et al. (2007***) have examined how bursting activity caused by retinal wave patterns can drive synaptic refinement of the RGC to LGN connections. They have developed a model for feedforward retinogeniculate development. Although these authors suggest that a novel learning rule is required for feedforward development, our model uses a Hebbian/anti-Hebbian learning rule that we believe is used throughout the brain and a complementary synaptogenic and shedding rule. Indeed we have used such synaptic modifications in a variety of adaptive synaptogenesis networks that include the visual system as well as in more abstract settings (***Levy and Colbert, 1991; Adelsberger-Mangan and Levy, 1993, 1994a***,b; ***Colbert et al., 1994; Thomas et al., 2015; Levy et al., 2016; Baxter and Levy, 2019, 2020***).

Studies of the development of RGC to SC, e.g. (***Savier et al., 2017***), support this supposition that feedforward development only requires Hebbian/anti-Hebbian modification.

In a recent paper (***Tikidji-Hamburyan et al., 2023***), the authors use a model of synaptic development that appears to be non-local and therefore non-biological. Nevertheless, they do point out the importance of convergence to stable connectivity which we agree is a sine qua non.

***Lee et al. (2013***) report a control group in which principal cell bursting is correlated with spindling. Their actual experimental group attempts to question the importance of such T-channel bursting in the generation of spindles. However, for these experimental animals that lack Cav3.1 (CACNA1G) there is no reason to believe that a compensatory Cav has taken the place of 3.1 since their mutated line of Cav knock-outs still develop spindling.

Concerning V1-L6 to LGN principal cell synaptogenesis, as far as we know, there are no direct, quantitative synaptic counts for the developing innervation. However, the anatomical data of ***Seabrook et al. (2013***) in mice can act as a proxy for such data, at least as an approximation. They quantify the fractional coverage of the LGN by labeled L6 axons. In their results (their Figure 4), postnatal day 4 (PN4) shows about 15% coverage of the LGN, at PN7 there is about 50% coverage, and at PN12, just before eye-opening, 100% coverage. Unfortunately, such a low-resolution technique cannot distinguish labeled axons from their labeled presynaptic structures. However, if we combine these anatomical observations with a series of electrophysiological measurements and functional manipulations, one becomes more comfortable with the idea that there is a large amount of feedback synaptogenesis occurring between PN8 and PN12. Indeed, this idea of axons invading in the vicinity of their targets prior to more precise activity-controlled synaptogenesis in modifying of existing synapses correlates with current theories of RGC to superior colliculus (SC) development (***Tsigankov and Koulakov, 2006***). For a similar general viewpoint in terms of feedforward development of the RGC to LGN, see ***Tikidji-Hamburyan et al. (2023***). Certainly, the one term that everyone agrees on is the ultimate determination of excitatory synapse formation, which is a Hebbian-like spike timing process (***Feldman, 2012; Markram et al., 2011***). Thus, in terms of biology, we assume that the labeling observed in ***Seabrook et al. (2013***) is initially dominated by axonal label with only a small fraction of the label due to presynaptic structures. As time progresses and synapses form, a larger fraction of the labeled area is due to label in presynaptic structures, and the retinotopy of connectivity begins rather crudely. But with time, PN9-PN12, and stage 3 retinal driving over this time period, a more precise point-wise retinotopy with smaller divergence (spread) occurs.

Motivating this interpretation that feedback synapses are developing rapidly from PN7 onward until eye-opening are the developmental electrophysiological data of ***Murata and Colonnese*** (***2016***). They provide evidence for the creation of functional feedback synapses, which follow upon the earlier development of feedforward, LGN to V1, excitatory connectivity. Like the RGC to SC retinotopy, the development of the L6 to LGN principal cell connectivity is dependent on spontaneous retinal activity (***Murata and Colonnese, 2016***). In the case of feedback to the LGN, it seems that the stage 3 retinal activity is more important than the stage 2 activity. ***Murata and Colonnese*** (***2016***) show that stronger spontaneous LGN firing occurs during the stage 3 period of retinal waves compared to smaller amounts of LGN bursting during the earlier stage 2 (their Fig 3). Moreover, (i) this strong, prolonged firing occurring in stage 3, disappears when V1 is functionally inactivated, and (ii) one of the spectral peaks of this strong, multi-unit firing occurs between 20 and 30 Hz, corresponding to the oscillation frequency of the developmental thalamocortical spindle event at this age. Consistent with this functional lesion result is their optogenetic activation of V1 (their Fig 4). This activation shows a much stronger response at PN10 compared to PN7. Thus, at PN7, V1 to LGN synapses exist, but at PN10, there are more of these synapses, or these synapses are stronger ones, or both. Interestingly, the frequency of the spindles themselves (these half-second or so events) occur in a burst-like manner, with many more spindle events at PN10 vs PN6. Thus, we conclude that there has been sparse synaptogenesis by PN7, producing the weak LGN responses to V1 activation, but by PN10, functional V1 to LGN synapses have proliferated substantially beyond the earlier connectivity of PN7. Moreover, the correlated increase of spindling events hints that the spindle events may be related to, or even the underlying cause of this synaptogenesis and synaptic strengthening.

### Enhancing the model

We claim there are straightforward biological hypotheses that convert our dynamic equations into dynamic biology. Moreover, our formulations are aimed at understanding Nature’s perspectives on producing appropriate developmental outcomes. The adaptive modification equations are written from two points of view (i) they are biologically implementable and (ii) they produce desirable representations and computations. Stability and useful computation are not the only constraints; it is also desirable that the algorithm minimizes both the time and energy devoted to development. This raises the question of the advantage of development over longer time scales. As can be seen in the simulations here, development in 80 passes takes about 25 minutes but *in vivo* development can last for three to five days. Thus, one wonders about the advantages of such slow development. One problem seems to be the maximal possible rate of axonal growth. A quick calculation, based on antler growth tells us that it takes 3.3 seconds for one micron of axonal growth. Perhaps this growth rate is limiting, but there may be some advantages in the slowness of such growth and the accuracy of weight convergence.

We have shown that random synaptogenesis can be restricted to speed up development and use less energy in development. We have listed five methods of restricting random synaptogenesis in Model, but we demonstrated only two of those methods here. While we have performed simulations using all five methods, the three methods not discussed here could be investigated further. Yet another possible mechanism to improve random synaptogenesis is to incorporate retrograde factors as illustrated in ***Baxter and Levy*** (***2020***) where a BDNF mechanism is proposed. Growing axons is expensive, but if the axons eligible for growth can be made more selective, energy will be saved.

We have not investigated the effect of synaptic failures on the development of feedback connections so that is another issue in need of research.

Here, we have assumed the feedforward connections are retinotopic and fixed. Thus, investigating the effect of a variety of learned feedforward (LGN to L6) synaptic weights, as well as the timing of the feedforward and feedback development, is another avenue of research.

### Suggested experimental studies

Here, we assume there is no memory or continuity of activity between spindle events because each spindle event occurs after the retinal input has shifted to a new retinotopic location. Thus, in the simulations, successive spindle events do not influence one another in terms of synaptic modification. Experiments are needed to confirm this non-interaction. Experiments are also needed to confirm that for any one spindle event the same principal LGN cells are re-firing.

In the simulations presented here, we have used a single LGN spike per burst within a spindle event instead of several high-frequency (140 to 250 Hz) spikes per burst. When multiple spikes per burst are specified, the development time tends to decrease. It would be useful to confirm this tendency experimentally.

Also, while we acknowledge that the microscopic experiments are quite difficult, actual synaptogenesis needs to be evaluated. This requires either super-resolution light microscopy or electron microscopy. Other difficult but important studies include the measurement of AMPA receptor to NMDA receptor ratios across the PN6 to eye-opening. The multi-unit activity studies (***Murata and Colonnese, 2016; Colonnese et al., 2017***) are helpful but actual single-unit recordings would be desirable. None of these studies are easy but all are technically feasible. In general, studies of synaptogenesis are difficult and inevitably require counting and count corrections with electron microscope resolution. An important technological challenge calls out for a marker of recent presynaptic growth. Perhaps someday there will be a way to label axons for recent actin polymerization and membrane insertion.

### Imprecision of reciprocal connectivity

A hallmark of several neural network theories is at the very least a degree of symmetric connectivity (***Hopfield, 1982; Cohen and Grossberg, 1983; Geman and Geman, 1984***). Such symmetry, for two neurons *j* and *k*, is expressed as *w*_*jk*_ = *w*_*kj*_. This symmetry assumption is at the heart of memory encodings that solve the pattern completion (filling-in) problem as in Hopfeld and Cohen-Grossberg networks and various random field models. In neuroscience, attractor networks, built by using such connectivity, are postulated to play an important role in the neural implementation of directed or focused attention (***Deco and Rolls, 2003***). Anatomical evidence for such connectivity exists for the feedback connections of L6 neurons to their primary exciting thalamic neurons (***Rockland, 2015***). Such anatomical results are a fundamental part of hodology in the hippocampal neocortical consolidation system (***Naber et al., 2001***).

In the simulations here, there is a propensity for the feedback synapses to spread wider than the retinotopic input to the feedforward-feedback circuit when the retinal inputs overlap spatially. It requires restrictions on synaptogenesis to get in-register, reciprocal connectivity with this minimal model consisting of a single set of dendritic weights per neuron when the active retinal inputs of two different spindle events overlap. However, it is quite possible that a less minimal model, perhaps consisting of neurons with multiple dendrites, could enhance the precision of the reciprocal connectivity. A more detailed modeling study is required to investigate this issue.

### From visual development to memory consolidation

Following feedforward retinogeniculate and thalamocortical development, spindling events are significantly enhanced and this correlates with synaptogenesis of the feedback corticothalamic synapses. Indeed, spindling may be a necessary condition for feedback synaptogenesis and synaptic modification. This hypothesis suggests a correlation between spindling and memory consolidation events. It is generally believed that sharp-wave ripple events coming out of the hippocampus lead to spindling in neocortex. We have previously argued (***Colbert et al., 1994***) that synaptogenesis is a necessary aspect of memory consolidation, and we are now beginning to consider a special role down to the biophysical level for the control of neocortical synaptogenesis by the hippocampus linked through neocortical spindling.

Although the modeling here concerns development between the LGN and V1, a related mechanism is likely used for memory consolidation between the hippocampus and neocortex, where there is a requirement for synaptogenesis. In fact, we translate the retinotopy alignment problem of visual feedback development into a problem to be solved as part of memory consolidation. In particular, there is a memory alignment problem that goes along with a theory of memory consolidation. It seems straightforward that the changing and novel representations that are part of adult consolidation must maintain a useful hippocampus. The hippocampus must be able to encode neocortical representations perhaps with a large amount of lossy compression and the outflow of the hippocampus that somehow points to a specific subset of neocortical neurons that need to be connected.

The hippocampus, specifically the Dentate Gyrus (DG) and Cornu Ammonis subfield 3 (CA3), is well known to play a major role in learning and memory. The DG-CA3 system, presumably having the responsibility of discovering unassociated (or miss-associated) aspects of experience, must be able to point to the neurons that should undergo synaptogenesis to encode this novel missed association. In order for the DG-CA3 system to remain useful throughout one’s lifetime, the mapping between the hippocampus and neocortex must frequently realign. We call this alignment between the hippocampus and neocortex “mnemotopy” which is analogous to retinotopy. However, retinotopy is stable, but mnemotopy is frequently changing. Indeed, the transformation between hippocampus and neocortex can be associated with memory consolidation, and, by hypothesis, could be the reason that spindling correlates with synaptogenesis as part of memory consolidation. The creation and maintenance of a memory topology implies a particular efficiency of memory storage.

## Conclusions

A minimal model of corticothalamic feedback featuring adaptive synaptogenesis including synaptic modification and shedding has been examined and demonstrated to produce nearly retinotopic and functionally useful connectivity from layer 6 of the visual cortex to the LGN. The model suggests why the T-channel and hyperpolarization are needed in the LGN and how these mechanisms provide spatially specific activities that lead to appropriate feedback connectivity.

The effects of varying the amount of RGC and LGN activity, the amount of L6 activity, and the LGN and L6 firing probabilities on the feedback synaptic weight development were examined and demonstrated. The total number of connections was demonstrated to be an approximately linear function of the number of active RGC neurons; similarly, the total number of connections was demonstrated to be an approximately linear function of the number of active L6 neurons. Increasing the number of active RGC cells was shown to increase the divergence of the feedback connections. Similarly, decreasing the number of active L6 neurons via lateral inhibition leads to decreased divergence but reduced synaptic strengths.

The adaptive synaptogenesis model includes synaptogenesis, synaptic modification, and synaptic shedding. Restricting synaptogenesis with presynaptic avidity and postsynaptic receptivity was shown to produce highly efficient synapse formation. While random synaptogenesis was shown to produce unstable connectivity, several methods of stabilizing connectivity and curtailing synaptogenesis were described. A weight sum thresholding method was demonstrated to be quite effective at stabilizing connectivity, ultimately developing approximately the same number of connections as restricted synaptogenesis. A spatial distance restriction on synaptogenesis that uses an attenuation function based on the distance between V1-L6 and LGN neurons was demonstrated to produce stable and retinotopic connectivity as long as the distance threshold is equal to or less than the number of active RGC neurons.

We hypothesize that developmental spindling is a result of selective principal cell activation by the retina upon the LGN, which leads to specific hyperpolarization of principal cells in the LGN. This cell-specific hyperpolarization not only halts the RGC-initiated firing, but activates T-channels in the dendrites of just those principal cells that were selectively fired. This selectivity is critical to the model as it creates a topographically specific memory of the retinal activation that is needed to establish retinotopic feedback connections from V1-L6 to LGN.

This study provides evidence that synaptogenesis is a necessary aspect of spindling and memory consolidation. We introduced the term “mnemotopy” to connote a memory topology. For example, the topological alignment, realignment, and refinement of synaptic connectivity between neural layers, and between hippocampus and neocortex, implies a memory topology. While retinotopy remains relatively stable after development, mnemtopy changes during memory consolidation.

## Acknowledgements

The authors are grateful to Estella Gaffney for providing many forms of assistance, including manuscript reviews and comments, during the preparation of this manuscript. The authors are also grateful to Thomas Charles Levy for support and encouragement during the preparation of this manuscript. The authors thank Dr. Costa Colbert for reviewing and providing comments on this manuscript. The authors also wish to thank Alexander Velikovich for help with tracking down and reviewing references. The authors thank Professor Mark Shaffrey for supporting this research. This research was funded in part by the Department of Neurosurgery in the School of Medicine at the University of Virginia, USA; and in part by Informed Simplifications, USA; and in part by Baxter Adaptive Systems, USA.

